# Inferring compound heterozygosity from large-scale exome sequencing data

**DOI:** 10.1101/2023.03.19.533370

**Authors:** Michael H. Guo, Laurent C. Francioli, Sarah L. Stenton, Julia K. Goodrich, Nicholas A. Watts, Moriel Singer-Berk, Emily Groopman, Philip W. Darnowsky, Matthew Solomonson, Samantha Baxter, gnomAD Project Consortium, Grace Tiao, Benjamin M. Neale, Joel N. Hirschhorn, Heidi L. Rehm, Mark J. Daly, Anne O’Donnell-Luria, Konrad J. Karczewski, Daniel G. MacArthur, Kaitlin E. Samocha

**Author notes:** These authors contributed equally: Michael H. Guo and Laurent C. Francioli. List of authors and their affiliations appear at the end of the paper. Correspondence should be addressed to K.E.S.

## Abstract

Recessive diseases arise when both the maternal and the paternal copies of a gene are impacted by a damaging genetic variant in the affected individual. When a patient carries two different potentially causal variants in a gene for a given disorder, accurate diagnosis requires determining that these two variants occur on different copies of the chromosome (i.e., are in *trans*) rather than on the same copy (i.e. in *cis*). However, current approaches for determining phase, beyond parental testing, are limited in clinical settings. We developed a strategy for inferring phase for rare variant pairs within genes, leveraging genotypes observed in exome sequencing data from the Genome Aggregation Database (gnomAD v2, n=125,748). When applied to trio data where phase can be determined by transmission, our approach estimates phase with 95.7% accuracy and remains accurate even for very rare variants (allele frequency < 1×10^−4^). We also correctly phase 95.9% of variant pairs in a set of 293 patients with Mendelian conditions carrying presumed causal compound heterozygous variants. We provide a public resource of phasing estimates from gnomAD, including phasing estimates for coding variants across the genome and counts per gene of rare variants in *trans*, that can aid interpretation of rare co-occurring variants in the context of recessive disease.

---

Determination of phase has important implications in clinical genetics, particularly in the diagnosis of recessive diseases that result from disruption of both copies of a gene. The disrupting bi-allelic variants can be either homozygous, where the same variant is present on both copies, or compound heterozygous, where two different variants are present on the two copies of the gene. Compound heterozygous variants present a challenge in genetic diagnosis because two variants observed within a gene in an individual can occur in *trans* or in *cis,* and only the former scenario results in compound heterozygosity. However, parental data are often not readily available for phasing or parents may not be available for follow-up testing, and short-read next generation sequencing largely cannot directly distinguish whether variant pairs are in *trans* or in *cis.* Thus, there is an important need for other approaches to accurately, easily, and cheaply determine phase of variant pairs.

The genetic relationship between a pair of variants on a haplotype can be disrupted by one of two processes: meiotic recombination and recurrent mutations. Meiotic recombination occurs more frequently in “hotspot” regions, and the probability of a recombination event occurring increases with distance between two variants^1^. A recurrent germline mutation event affecting a variant on a haplotype can also disrupt the genetic relationship of the variants on the haplotype. Rates of recurrent mutations are dependent on mutation type (e.g., transition versus transversion) and epigenetic marks (particularly CpG methylation), among other factors^2–6^. Thus, the rates of both meiotic recombination and mutation have important implications in determining the phase of variants.

There are several approaches for directly inferring phase for variant pairs observed in an individual. Phase may be determined directly using data from sequencing reads. However, for data from typical short-read sequencing technologies such as Illumina, read-based phasing methods are generally only possible for variants in close proximity to each other^7^, although some variant pairs at longer distances can be phased with more sophisticated algorithms^8–10^. Long-read sequencing technologies allow for direct determination of phase for variant pairs at longer distances, but these technologies are more expensive and have not yet been widely applied in clinical settings^11,12^. There are also laboratory-based molecular methods for determining phase of variant pairs, but these methods are low-throughput and technically challenging^13^. While phase can be determined based on transmission of variants from parents to offspring, this approach increases cost and may pose other logistical and ethical challenges, and is not always an option if parents are deceased, unavailable (e.g., living far away or incarcerated), unknown in the case of adoption, or unwilling to participate. These direct phasing approaches all thus present critical limitations for determining phase of variant pairs within an individual in a clinical setting.

Alternative, indirect approaches for phasing rely on statistical methods applied to population data (reviewed in Tewhey et al.^14^ and Browning and Browning^15^). Many of these approaches build off of the Li-Stephens model^16^ and leverage genetic data from large numbers of unrelated or distantly related individuals to identify shared haplotypes among individuals in a population^17–19^. However, these methods require a large number of reference samples (typically n ∼10^5^-10^6^ individuals) and are computationally intensive to perform. These approaches perform less well for rare variants. Furthermore, these approaches cannot be readily applied to exome sequencing data which does not provide enough density of surrounding variants to allow for accurate phasing. Despite these limitations, these population-based approaches are attractive because they do not require sequencing of additional family members or application of expensive sequencing approaches.

In this work, we sought to address existing challenges of phasing in clinical settings, particularly with regard to rare variants observed in exome sequencing data. We implement an approach that leverages the principles of population-based phasing by estimating haplotype patterns from a large reference population and using these patterns to infer variant phase in an individual. We cataloged the haplotype patterns of rare coding and flanking intronic/UTR variants within genes using the Genome Aggregation Database (gnomAD), which performed aggregation and joint genotyping of exome sequencing data from 125,748 individuals^20^. We then demonstrate that we can leverage these data to generate a resource for phasing rare coding variants observed in an individual, and identify factors that influence the accuracy of our approach. Additionally, we provide statistics for how often different types of variants are observed in *trans* within gnomAD, stratified by AF and mutational consequence, to provide a background rate contextualization when observing biallelic rare variants in rare disease cases. Finally, we disseminate these resources in a user-friendly fashion via the gnomAD browser for community use.

## Results

### Inference of phase in gnomAD

We sought to address the challenges of phasing variants observed in individual samples in clinical settings by applying the principles of population-based phasing. Specifically, to infer the phase of variants in an individual, we leveraged the fact that haplotypes are usually shared across individuals in a population (**Fig. 1a**). If two variants are in *cis* in many individuals in a population, then they are likely to be *cis* in any given individual’s genome. Similarly, if two variants are in *trans* in other individuals in a population, then they are likely to be *trans* in any given individual’s genome. This latter scenario also provides information that the variant combination may be tolerated in *trans* since it has been found in an individual in gnomAD. We reasoned that by generating phasing estimates from a large reference population, we could infer the phase of variants observed in an individual.

**Fig. 1:**
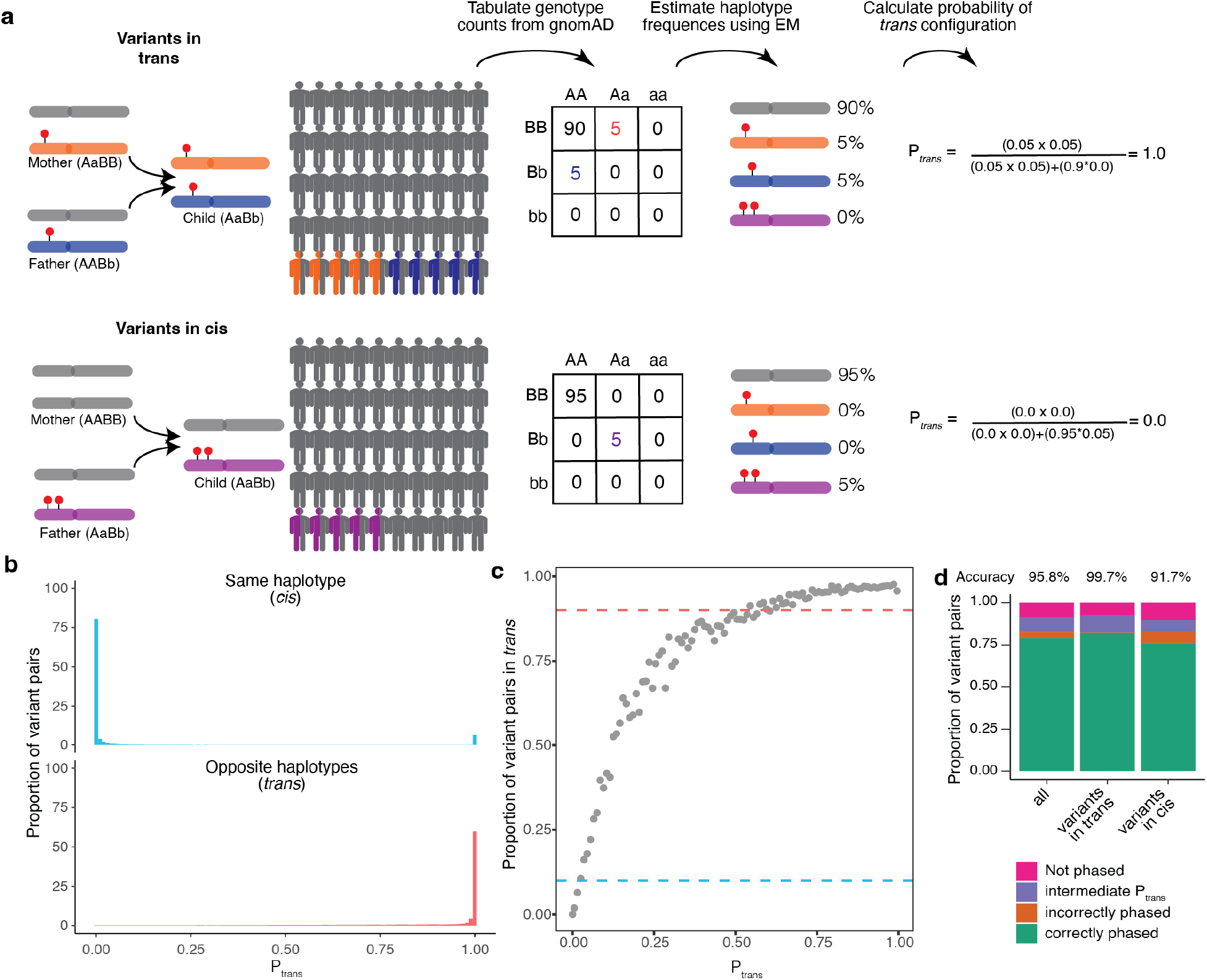
Overview of phasing approach using Expectation-Maximization method in gnomAD. **a,** Schematic of phasing approach. **b,** Histogram of P_trans_ scores for variant pairs in *cis* (top, blue) and in *trans* (bottom, red). **c,** Proportion of variant pairs in each P_trans_ bin that are in *trans*. Each point represents variant pairs with P_trans_ bin size of 0.01. Blue dashed line at 10% indicates the P_trans_ threshold at which ≥ 90% of variant pairs in bin are on the same haplotype (P_trans_ ≤ 0.02). Red dashed line at 90% indicates the P_trans_ threshold at which ≥ 90% of variant pairs in bin are on opposite haplotypes (P_trans_ ≥ 0.55). Calculations are performed using variant pairs with population AF ≥ 1×10^−4^. **d,** Performance of P_trans_ for distinguishing variant pairs in *cis* and *trans.* Accuracy is calculated as the proportion of variant pairs correctly phased (green bars) divided by the proportion of variant pairs phased using P_trans_ (orange plus green bars). **b-d,** P_trans_ scores are population-specific.

Predicting the phase of a given pair of variants in an individual first requires that we estimate the haplotype frequencies in the population for a given pair of variants. To estimate haplotype frequencies, we used exome sequencing samples from gnomAD v2, a large sequencing aggregation database^20^. In total, there were 125,748 exome sequencing samples after rigorous sample and variant quality control (Online Methods). There are several key advantages of using gnomAD as a reference dataset for calculating haplotype frequencies. First, samples in gnomAD undergo uniform processing and variant-calling, mitigating the impact of technical artifacts. Second, with over 125,000 individuals in gnomAD, the database provides sufficient sample sizes to estimate haplotype frequencies below 1×10^−5^. Lastly, gnomAD offers significant genetic ancestral diversity, allowing results of our study to be applied beyond samples with European ancestry.

We focus in this study on pairs of rare exonic variants occurring in the same gene, which are of the greatest interest in the context of Mendelian conditions. We required both variants to have a global minor allele frequency (AF) in gnomAD exomes <5% and required variants to be coding, flanking intronic (from position −1 to −3 in acceptor sites, and +1 to +8 in donor sites) or in the 5’/3’ UTRs. This encompassed 5,320,037,963 unique variant pair combinations across 19,877 genes. Of these variant pairs, 11,786,014 are carried by the same individual at least once in gnomAD, of which only 105,322 are both singleton variants and seen in the same individual, where we are unable to make a phase prediction. We performed estimates based on all exome sequencing samples in gnomAD v2, as well as separate estimates within each genetic ancestry group (African/African American [AFR]: n=8128; Admixed American [AMR]: 17296; Ashkenazi Jewish [ASJ]: 5040; East Asian [EAS]: 9179; Finnish [FIN]: 10824; non-Finnish European [NFE]: 56885; Remaining: 3070; South Asian [SAS]: 15308).

For each pair of variants, we first generated pairwise genotype counts in gnomAD, with nine possible pairwise genotypes for each pair of variants (**Fig. 1a**). We then applied the Expectation-Maximization (EM) algorithm to each pair of variants to generate haplotype frequency estimates based on the observed pairwise genotype counts^21^. For a given pair of variants observed in an individual, the probability of two variants being in *trans* (P_trans_) is the probability of inheriting each of the haplotypes that contain only one of the two variant alleles.

### Validation of phasing estimates using trio data

To measure the accuracy of our approach, we analyzed variants in a set of 4,992 trios that underwent exome sequencing and joint processing with gnomAD. In this trio structure, we could accurately measure phase using parental transmission as a gold standard and could compare with phase as predicted using the EM algorithm in gnomAD samples. We first estimated the genetic ancestry of each individual in the trios by projecting on the principal components of ancestry in the gnomAD v2 samples (**Supplementary Fig. 1**). Of the 4,992 children from the trios, 4,775 were assigned to one of seven genetic ancestry groups (AFR: 73; AMR: 358; ASJ: 62; EAS: 1252; FIN: 149; NFE: 2815; SAS: 46). For validating and measuring accuracy of our approach, we removed from gnomAD any samples in our trio dataset that did not fall into one of the seven aforementioned genetic ancestry groups. We used our method leveraging gnomAD data to estimate phase for every pair of rare (global AF < 5% and population AF < 5%) coding and flanking intronic/UTR variants within genes observed in either of the parents. Across the 4,775 trio samples, we identified 339,857 unique variant pairs and 1,115,347 total variant pairs (mean 241.7 variant pairs per trio sample) (**Supplementary Fig. 2a**). On average, each trio sample had 64.4 variant pairs where both variants were missense, inframe insertions/deletions (indels) or predicted loss-of-function (pLoF), and 0.35 pLoF/pLoF variant pairs (**Supplementary Fig. 2b-c**). Nearly all of the variants identified in the trios were single nucleotide variants, with only 2.7% being short indels. A breakdown of functional consequences for these variants is depicted in **Supplementary Fig. 3a**.

The vast majority (91.1%) of unique variant pairs seen in the trio samples were observed in gnomAD at least once and thus amenable to our phasing approach (**Fig. 1d**). By contrast, only 2.1% of variant pairs in these samples were within 10 bp of each other, the range in which we previously found read-back phasing of the physical read data to be most effective^7^ (**Supplementary Fig. 3b**). Additionally, we find that 8.2% of variant pairs were within 150 bp, the typical length of an Illumina exome sequencing read 19.2% of variant pairs were within the same exon. Thus, our approach has a much higher ability to phase variants than physical read-back phasing data.

For each variant pair, we calculated the probability of being in *trans* (P_trans_) based on the haplotype frequency estimates in gnomAD as described above. We found a bimodal distribution of P_trans_ scores; that is, the majority of probabilities were either very high (> 0.99; suggesting a high likelihood of being in *trans*), or they were very low (< 0.01; suggesting a high likelihood of being in *cis*) (**Fig. 1b, Supplementary Fig. 4a-g**). Using the trio phasing-by-transmission data as a gold standard, we generated receiver-operator curves for distinguishing whether a variant pair is likely in *trans* and found that our approach achieved high sensitivity and specificity (area under curve [AUC] ranging from 0.892 to 0.997 across the component genetic ancestry groups) (**Supplementary Fig. 5a**) and high precision and recall (**Supplementary Fig. 5b**).

We next defined P_trans_ thresholds for classifying variants as being in *cis* versus *trans* (see Methods for additional details). To set these thresholds, we first binned variant pairs observed in the trio data based on their P_trans_ scores calculated from gnomAD samples from the same genetic ancestry group. We used only variants on odd chromosomes (i.e., chromosomes 1, 3, 5, etc) to determine P_trans_ thresholds. For each P_trans_ bin, we calculated the proportion of trio variant pairs that were in *cis* or *trans* based on trio phasing-by-transmission. The P_trans_ threshold for variant pairs in *trans* was defined as the minimum P_trans_ such that ≥ 90% of variant pairs in that bin were in *trans* based on trio phasing-by-transmission. Similarly, the P_trans_ threshold for variants in *cis* was defined as the maximum P_trans_ such that ≥ 90% of variant pairs in that bin were in *cis* based on trio phasing by transmission. This resulted in P_trans_ thresholds of ≤ 0.02 and ≥ 0.55 as the threshold for variants in *cis* and *trans*, respectively (**Fig. 1c**).

We next assessed how well our P_trans_ thresholds performed by measuring phasing accuracy using the EM algorithm against trio phasing-by-transmission as a gold standard. For measuring accuracy, we utilized only variant pairs observed on even chromosomes (i.e., chromosomes 2, 4, 6, etc). Overall, 91.1% of unique variant pairs had both variants present in the corresponding population in gnomAD and therefore were amenable to phasing (**Fig. 1d**), with only a minority (8.6%) of unique variant pairs having an intermediate P_trans_ score (i.e., 0.02 < P_trans_ < 0.55) and thus an indeterminate phase. We calculated accuracy as the percentage of phaseable variant pairs (i.e., both variants present in gnomAD, and P_trans_ score ≤ 0.02 or ≥ 0.55) that were correctly phased. Based on these P_trans_ thresholds, the overall phasing accuracy was 95.8%. The accuracy for unique variant pairs that are in *cis* based on trio data was 91.7%, and the accuracy for variant pairs in *trans* was 99.7%.

We also calculated the overall percentage of variants correctly phased in a given individual (i.e., variants are counted more than once if seen multiple times in the trio data). 96.9% variant pairs in a given individual had both variants present in gnomAD and therefore were amenable to phasing, and 92.3% of variant pairs observed in a given individual were correctly phased using our approach. Accuracy is lower, but still high, for rare variants; notably, among variant pairs with AF < 0.1%, 80.1% of variant pairs in a given individual were correctly phased. Together, these results suggest that our approach can generate highly accurate phasing estimates.

### Accuracy of phasing across allele frequencies

Since the variants that are most likely to be of interest in clinical genetics are rare, we assessed the accuracy of phasing at different AF bins. We found high accuracy (i.e., proportion correct classifications) ranging from 0.779 to 0.988 across pairs of AF bins (**Fig. 2**). In general, accuracy remained high across allele frequencies for variant pairs in *trans* based on trio phasing data. For variant pairs in *cis* based on trio phasing data, accuracy was high when the allele frequencies of both variants in the pair were high (AF ≥ 0.001). However, accuracy was much lower for rare variants in *cis* (AF < 1×10^−4^), and in particular when one variant in the pair is rare and the other is more common (**Fig. 2c**). Variant pairs where both variants are singletons (i.e., observed once in gnomAD) were phased well for variants in *trans* based on the trio phasing data (accuracy of 0.993). Given the lack of information, we do not report the phasing estimates for singleton/singleton variant pairs in *cis* (see Discussion).

**Fig. 2:**
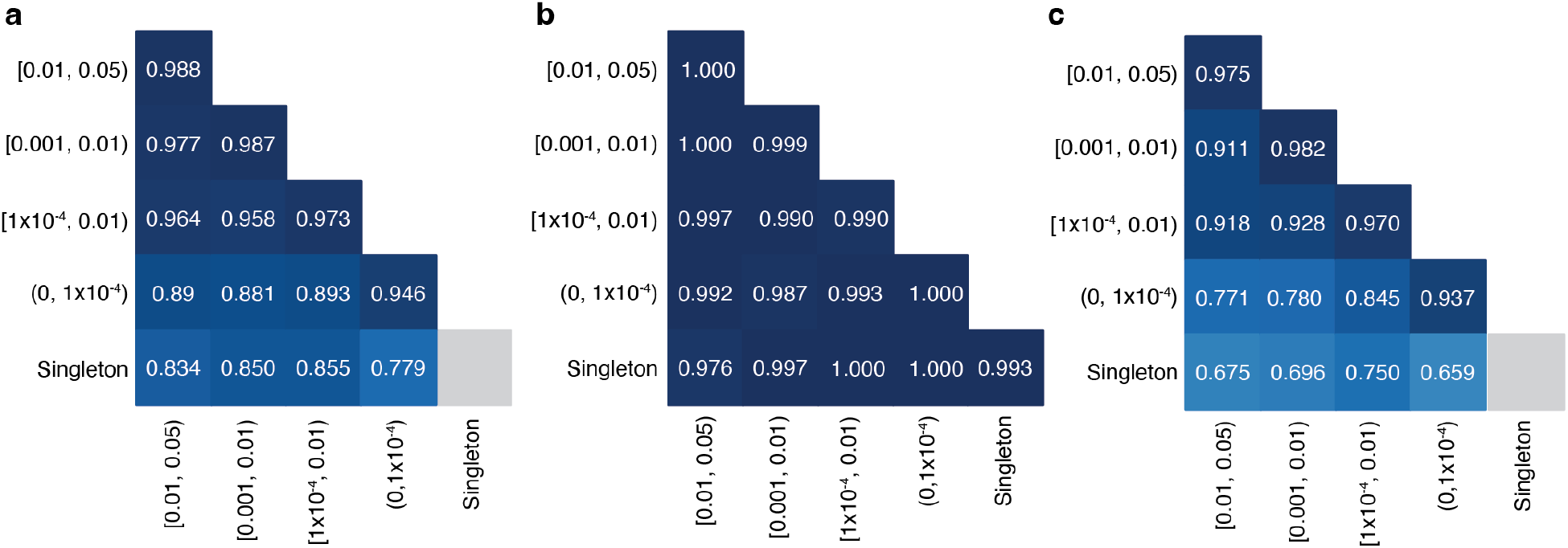
Phasing accuracy as a function of variant allele frequency (AF). Phasing accuracy at different AF bins for all variant pairs (**a)**, variant pairs in *trans* (**b)**, and variant pairs in *cis* (**c)**. Shading of squares and numbers in each square represent the phasing accuracy. Y-axis labels refer to the more frequent variant in each variant pair and X-axis labels refer to the rarer variant in each variant pair. Accuracy is the proportion of correct classifications (i.e., correct classifications / all classifications) and is calculated for all unique variant pairs seen in the trio data across all populations using population-specific P_trans_ calculations.

### Accuracy of phasing across genetic ancestry groups

In the above analyses, we used P_trans_ estimates calculated from samples in gnomAD with the same genetic ancestry group (“population”) in which the variant pair was seen in the trio data. We next asked if using all samples in gnomAD to calculate P_trans_ (“cosmopolitan”) would improve accuracy given larger sample sizes from which to calculate P_trans_ (**Supplementary Fig. 6**), with the caveat that using the full set of gnomAD samples would result in some genetic ancestry mismatching. Across populations, we found that accuracy was generally similar when using population-specific ancestry estimates as compared to cosmopolitan estimates (**Fig. 3a-b**). However, for certain genetic ancestry groups such as AFR and EAS, accuracy was slightly lower when using cosmopolitan estimates as compared to population-specific estimates specifically for variants in *trans* in these populations. For example, the phasing accuracy for variants in *trans* in the AFR ancestry group was 0.995 when using AFR-specific P_trans_ estimates, but drops to 0.952 when cosmopolitan P_trans_ estimates are used. These results suggest that cosmopolitan estimates allow a greater proportion of variants to be phased with generally similar accuracy as population-specific estimates, though we do identify certain scenarios where more caution is required.

**Fig. 3:**
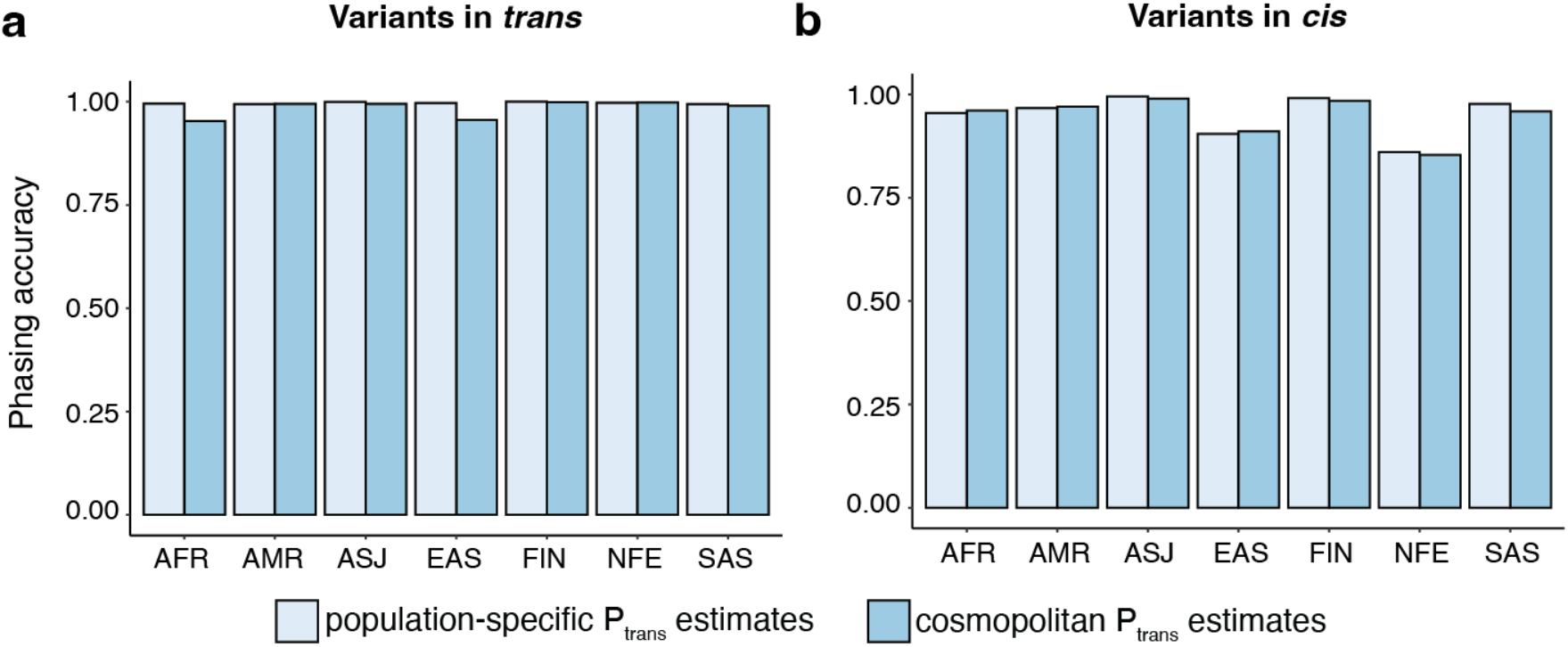
Phasing accuracy using population-specific versus cosmopolitan P_trans_ estimates. Population-specific P_trans_ estimates are shown in light blue and cosmopolitan P_trans_ estimates are shown in medium blue. Accuracies are shown separately for variants in *trans* (**a**, left) and variants in *cis* (**b**, right)

### Effect of variant distance and mutation rates on phasing accuracy

One factor that is predicted to affect the accuracy of our phasing estimates is the frequency of meiotic recombination between pairs of variants, as recombination events would disrupt the haplotype configuration of the variant pairs. To further explore the impact of recombination, we plotted the accuracy of our phasing estimates as a function of physical distance between variant pairs. We found that for variants in *trans,* the accuracy of phasing was maintained across physical distances. However, for variant pairs in *cis,* the accuracy rapidly decreased with longer physical distances (**Fig. 4a**). Since physical distance is only a proxy for recombination frequency, we also performed this analysis using interpolated genetic distances (**Fig. 4b**). We found again that variants in *trans* had preserved phasing accuracy across genetic distances, while variants in *cis* had phasing estimates that decreased substantially with genetic distance, particularly at distances greater than 0.1 centiMorgan.

**Fig. 4:**
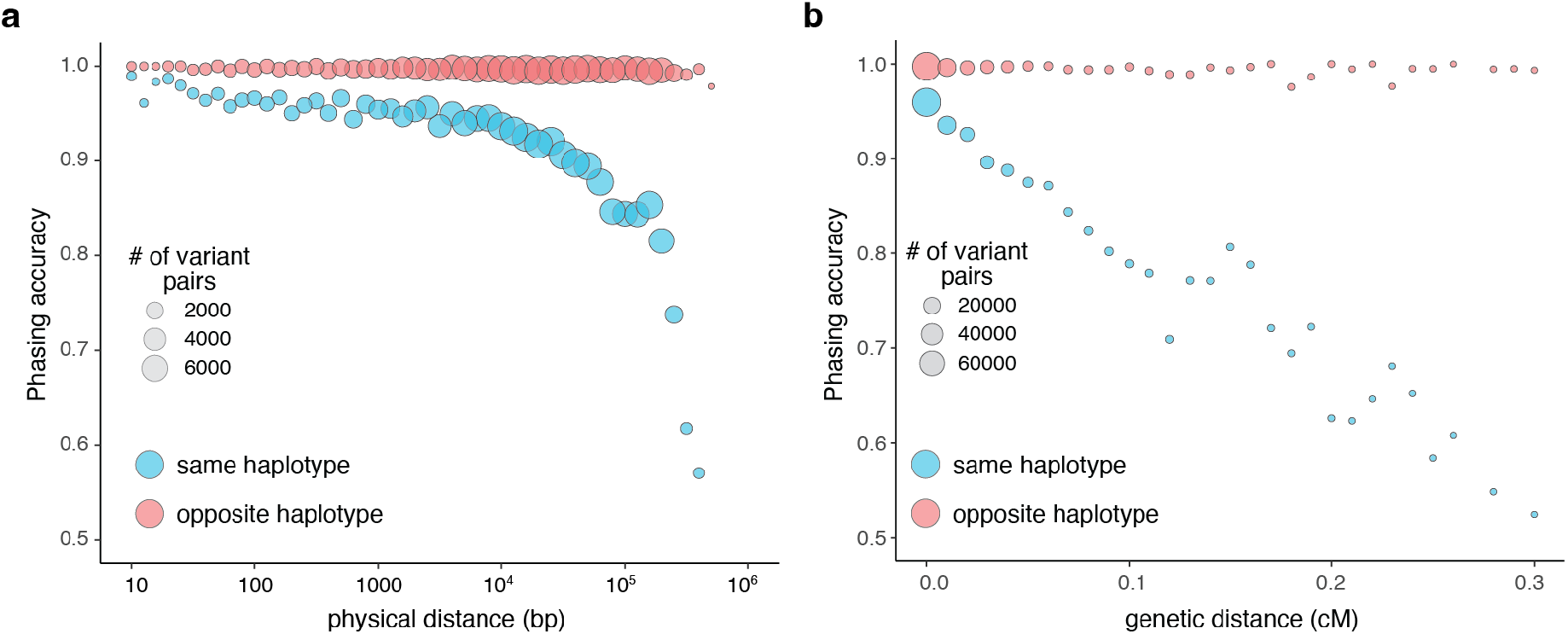
Phasing accuracy as a function of distance between variant pairs. **a,** Phasing accuracy (y-axis) as a function of physical distance (in base pairs on log_10_ scale) between variants (x-axis). Blue represents variants on the same haplotype (in *cis*), and red represents variants on opposite haplotypes (in *trans*). **b,** Same as **a,** except the x-axis shows genetic distance (in centiMorgans). Accuracies for **a** and **b** are based on unique variant pairs observed across all genetic ancestry groups and are calculated using population-specific P_trans_ estimates.

We also tested the effect of recombination by examining a set of 20,319 multinucleotide variants (MNVs), which are pairs of genetic variants in *cis* that are very close together in physical distance (≤ 2 bp) and thus have minimal opportunity for recombination between them. These variants have previously been accurately phased using physical read data^22,23^. When examining this set of MNVs, we found that the phasing accuracy using our approach was 96.0%, with only 3.5% of MNVs phased incorrectly (the remaining 0.46% had indeterminate phasing estimates). We found that the vast majority of MNVs that were phased incorrectly using P_trans_ had disparate AFs between the two variants, similar to what we had observed above (data not shown).

Another potential driver of inaccurate phasing is recurrent germline mutations, and the rate at which recurrent germline mutations occur can be highly variable. Transitions have higher mutation rates than transversions^2,24^. Furthermore, CpG transitions have the highest mutation rates among single nucleotide changes, with mutation rates increasing with higher methylation rates at the CpG sites^20^. To better understand the impact of mutation rates on phasing accuracy, we classified each single nucleotide variant (SNV) in the trio data as a transversion, non-CpG transition, or CpG transition. For CpG transitions, we further classified the SNV as having low, medium, or high DNA methylation as before^20^. We then calculated phasing accuracy as a function of combinations of mutation types using the trio data (**Supplementary Fig. 7a-c**). We found similar accuracy for transversions and transitions (∼0.97) (**Supplementary Fig. 7a)**. However, we found that the mutation rates had a strong impact on variant pairs in *cis* but not those in *trans* (**Supplementary Fig. 7b-c**). For variants in *cis,* the phasing accuracies were lower at medium and high methylation CpG sites (0.82-0.89) than they were for low methylation sites (0.96). These results are consistent with recurrent mutations contributing to inaccurate phasing estimates, particularly for variant pairs in *cis*.

### Evaluation of limitations of phasing approach

We next sought to address several limitations we observed in our current phasing data. First, 4.7% of variant pairs in the 4,775 trios were not present in gnomAD and thus not amenable to phasing even using cosmopolitan P_trans_ estimates. To understand how the proportion of variants amenable to phasing changes as a function of gnomAD reference sample size, we performed a subsampling analysis of gnomAD from 121,912 (all of gnomAD v2 after removing overlapping trio samples) down to 1,000, 10,000, or 100,000 samples (**Supplementary Fig. 8a**). We found that subsampling greatly reduces the proportion of variants amenable to phasing, but that accuracy is generally preserved. For example, when subsampling down to 10,000 samples, just 76.4% of variant pairs observed in the trios were amenable to phasing, but phasing accuracy remained high (91.9%) when using cosmopolitan P_trans_ estimates (compared to 93.6% accuracy when using the full gnomAD cohort).

We next assessed variant pairs with intermediate EM scores (0.02 < P_trans_ < 0.55) where our approach gives an indeterminate phase estimate. We found that nearly all (99.8%) of the variant pairs with intermediate P_trans_ scores included more common variants (AF ≥ 0.001) (**Supplementary Fig. 8b**). For variant pairs where the more common variant had AF ≥ 0.001, 9.5% of variant pairs had an intermediate P_trans_ score. In contrast, variant pairs where the more common variant with AF < 0.001, just 0.19% of variant pairs had an intermediate P_trans_ score. Intermediate P_trans_ scores can only occur when all four haplotypes are observed. For rare variants, typically not all four haplotype combinations due to sampling and because rare variants are younger and have less opportunity for recombination/recurrent mutation to generate all haplotype combinations.

Finally, we investigated the seemingly counterintuitive observation that phasing accuracy was lowest for NFE, where we had the highest number of gnomAD reference samples. We postulated that this apparent lower phasing accuracy in NFEs might be due to the larger number of trios we tested (for example, we tested 2815 NFE trio samples compared to 73 AFR samples), rather than an issue with phasing of NFE samples in the gnomAD reference dataset itself. To test this, we randomly subsampled the NFE trio samples from 2815 trios down to 282 (10%), 563 (20%) or 1408 (50%) trios. Upon subsampling, we found that a smaller proportion of unique variant pairs were in lower AF bins (< 1×10^−4^) where phasing is most challenging (**Supplementary Fig. 8c**), with a corresponding improvement in accuracy upon subsampling of the trio samples (**Supplementary Fig. 8d**). These results suggest that the observation of a lower phasing accuracy in NFE is an artifact of ascertaining and testing a larger number of NFE trio samples. Intuitively, this artifact results from our approach of measuring accuracy using unique variant pairs within a population. With increasing numbers of trios tested in our trio validation set, more common variant pairs where phasing accuracy is higher are observed multiple times yet counted only once. In contrast, with larger numbers of trios tested, we observe a larger number of unique rare variant pairs where accuracy is lower.

### Demonstration of accuracy in a cohort of patients with Mendelian disorders

To demonstrate our approach in a clinically relevant situation, we turned to a set of 627 patients from the Broad Institute Center for Mendelian Genetics (CMG)^25^. All patients had a confident or strong candidate genetic diagnosis of a Mendelian condition based on carrying two rare variants in a recessive disease gene consistent with the patient’s phenotype. Across the 627 diagnoses, we were able to estimate phase for the 293 patients for which both variants were present in gnomAD (**Supplementary Table 1**). For the analysis of these 293 variant pairs, we used the population-specific P_trans_ estimate when available (n=215), and when not, we used the cosmopolitan P_trans_ estimates (n=78). Our phasing approach indicated 281 (95.9%) variant pairs to be in *trans*, seven variant pairs (2.4%) to be in *cis*, and five (1.7%) as indeterminate (0.02 < P_trans_ < 0.55 or singleton-singleton variant in the same individual). Had only the cosmopolitan P_trans_ estimates been used, one of the 281 in *trans* predictions would have been predicted in *cis* and one indeterminate. Of the seven variant pairs indicated to be in *cis*, six originated from patients with proband-only sequencing. For these patients, the responsible clinician was contacted to ensure phenotype overlap with the disease gene and to pursue parental Sanger sequencing for confirmatory phasing by transmission or long read sequencing, where possible. The remaining variant pair predicted to be in *cis* originated from a patient with trio-sequencing, which confirmed the variants to be in *trans*, and our inferred phase to be incorrect. Further details on the incorrect prediction are provided in **Supplementary Table 1**. Despite one error and several indeterminate predictions, overall, the results suggest that our phasing approach is highly accurate in clinical scenarios in patients with suspected Mendelian conditions and can be informative for a large fraction (just under 50% in our cohort) of candidate diagnoses, dependant on presence of both variants in gnomAD.

### Bi-allelic predicted damaging variants

Next, we tabulated for each gene the number of individuals in gnomAD who carry two rare heterozygous variants, stratified by the predicted phase using P_trans_ cutoffs (i.e., in *trans,* unphased [intermediate P_trans_], and in *cis*), AF, and the predicted functional consequence of the least damaging variant in the pair. For comparison, we tabulated individuals with homozygous variants in the same manner. We classified predicted functional consequences as pLoF, missense with deleteriousness scored by REVEL^26^ in line with recent ClinGen recommendations^27^, and synonymous. These data are available for gnomAD v2 as a downloadable table with the counts of individuals by phase for each gene across a total of 26 consequences and five AF thresholds.

Overall, the number of individuals with rare, compound heterozygous (in *trans*), predicted damaging variants was low (median 0 individuals per gene with compound heterozygous loss-of-function variants at ≤ 1% AF, range 0-9) and only occurred in a small number of genes (**Fig. 5** and **Supplementary Fig. 9**). Twenty eight genes carried compound heterozygous pLoF variants (in 56 individuals) and an additional four genes carried compound heterozygous variants with at least a strong REVEL missense predicted consequence (in six individuals) at ≤ 1% AF cutoff. The vast majority of these genes have not, to date, been associated with disease (**Fig. 5b**). Manual curation of the pLoF variants resulted in seven high confidence “human knock-out” genes (*ARHGEF37, CCDC66, FAM81B, FYB2, GNLY, RBKS,* and *SDSL*). These genes are not associated with Mendelian disease nor are they known to be essential (see Methods for additional details). In the remaining 21 of the 28 genes with compound heterozygous pLoF variants, true loss-of-function was found to be uncertain or unlikely following manual curation, due, for example, to the variant falling in the last exon of the gene, in a weakly conserved exon, or in the minority of transcripts (**Supplemental Table 2**). We previously manually curated all homozygous pLoF variants in gnomAD^20^. The absence of rare compound heterozygous “human knock-out” events in essential genes was expected given that gnomAD is largely depleted of individuals with severe, early-onset conditions.

**Fig. 5:**
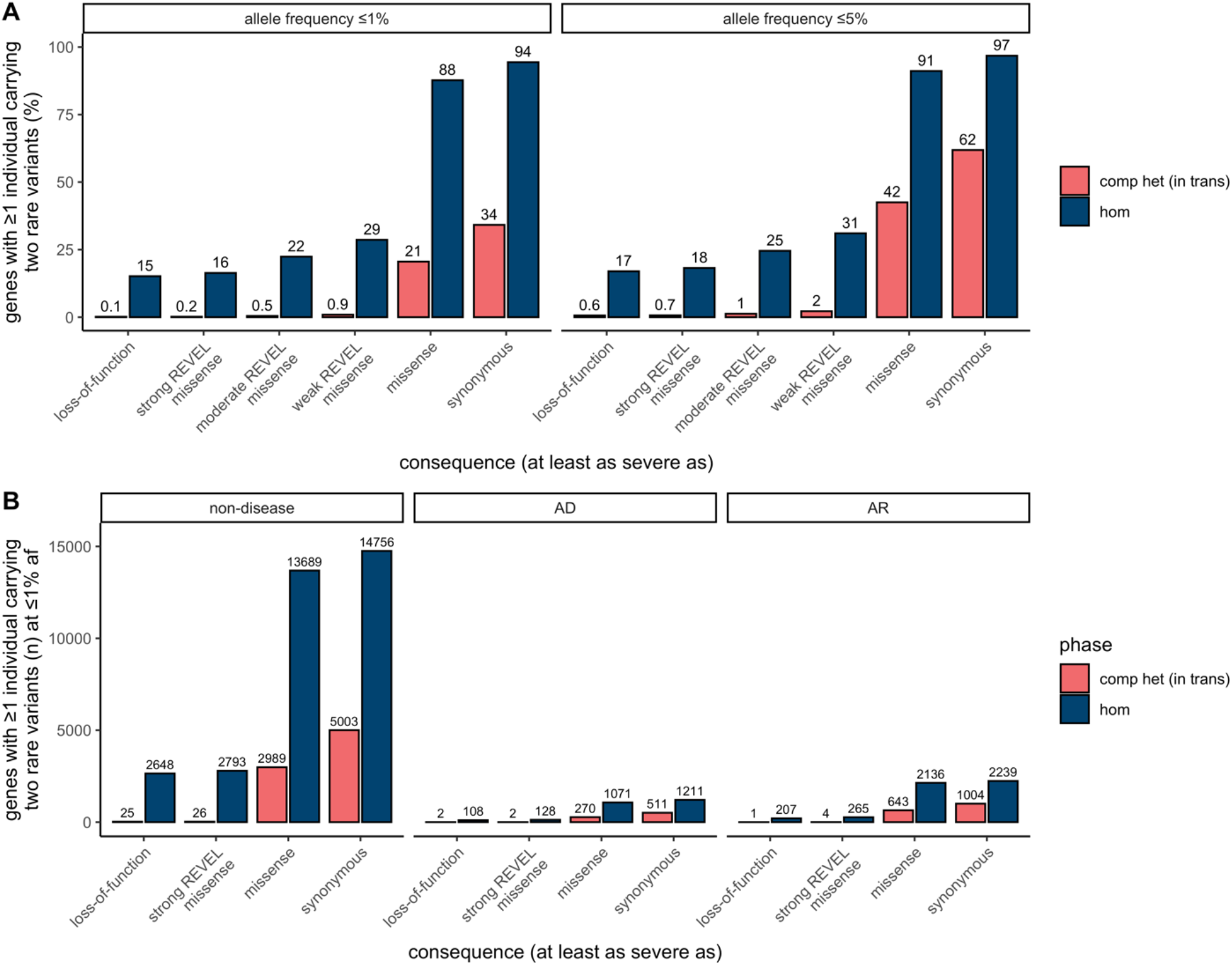
Counts of genes with variants in *trans* in gnomAD. **a,** Proportion of genes with one or more individuals in gnomAD carrying predicted compound heterozygous (in *trans*) variants or a homozygous variant at ≤ 1% and ≤ 5% AF stratified by predicted functional consequence. **b,** Number of genes with ≥ 1 individual in gnomAD carrying compound heterozygous (in *trans*) or homozygous predicted damaging variants at ≤ 1% AF, stratified by predicted functional consequence and Mendelian disease-association in the Online Mendelian Inheritance in Man database (OMIM). In total, 28 genes (25 non-disease, 2 AD, and 1 AR) carried predicted compound heterozygous loss-of-function variants at ≤ 1% AF, only seven of which were high confidence “human knock-out” events following manual curation. For predicted compound heterozygous variants, both variants in the variant pair must be annotated with a consequence at least as severe as the consequence listed (i.e., a compound heterozygous loss-of-function variant would be counted under the pLoF category but also included with a less deleterious variant under the other categories). All homozygous pLoF variants previously underwent manual curation as part of Karczewski et al^20^. AF, allele frequency; comp het, compound heterozygous; hom, homozygous; AD, autosomal dominant; AR, autosomal recessive.

### Generation of public resource

To make our resource widely usable to both clinicians and researchers, we have calculated and released pairwise genotype counts and phasing estimates for each pair of rare coding variants occurring in the same gene for gnomAD. These genotype counts are phasing estimates shown for all pairs of variants within a gene where both variants have global minor AF (or population-specific frequency) in gnomAD exomes < 5%, and are either coding, flanking intronic (from position −1 to −3 in acceptor sites, and +1 to +8 in donor sites) or in the 5’/3’ UTRs. We have integrated these data into the gnomAD browser so that clinicians and researchers can easily look up a pair of variants to obtain the genotype counts, haplotype frequency estimates, P_trans_ estimates, and likely co-occurrence pattern (**Fig. 6a**). These results are shown for each individual genetic ancestry group and across all genetic ancestries in gnomAD v2. In addition, the data are available as a downloadable table for all variant pairs that were seen in at least one individual.

**Fig. 6:**
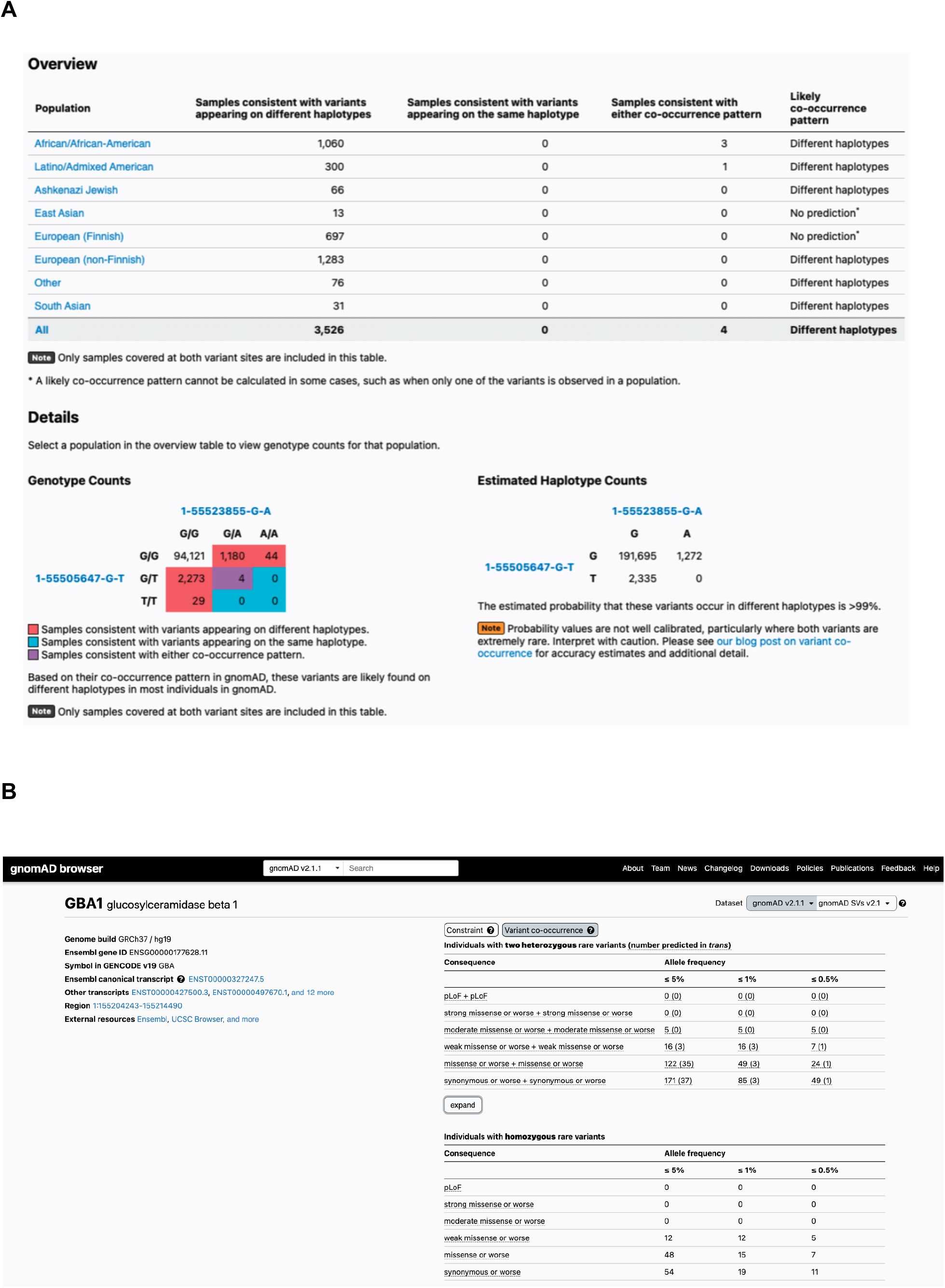
Publicly-available browser for sharing of phasing data. **a,** Sample gnomAD browser output for two variants (1-55505647-G-T and 1-55523855-G-A) in the gene *PCSK9*. On the top, a table subdivided by genetic ancestry group displays how many individuals in gnomAD from that genetic ancestry are consistent with the two variants occurring on different haplotypes (*trans*), and how many individuals are consistent with their occurring on the same haplotype (*cis*). Below that, there is a 3×3 table that contains the 9 possible combinations of genotypes for the two variants of interest. The number of individuals in gnomAD that fall in each of these combinations are shown and are colored by whether they are consistent with variants falling on different haplotypes (red) or the same haplotype (blue), or whether they are indeterminate (purple). The estimated haplotype counts for the four possible haplotypes for the two variants as calculated by the EM algorithm is displayed on the bottom right. The probability of being in *trans* for this particular pair of variants is > 99%. **b,** Variant co-occurrence tables on the gene landing page. For each gene (*GBA1* shown), the top table lists the number of individuals carrying pairs of rare heterozygous variants by inferred phase, AF, and predicted functional consequence. The number of individuals with homozygous variants are tabulated in the same manner and presented as a comparison below. AF thresholds of ≤ 5%, ≤ 1%, and ≤ 0.5% are displayed across six predicted functional consequences (combinations of pLoF, various evidence strengths of predicted pathogenicity for missense variants, and synonymous variants). Both variants in the variant pair must be annotated with a consequence at least as severe as the consequence listed (i.e., pLoF + strong missense also includes pLoF + pLoF).

Furthermore, on the landing page of each gene in the gnomAD v2 browser, we have incorporated counts tables detailing the number of individuals carrying two rare variants stratified by AF, and functional consequence. The first table counts individuals carrying two rare heterozygous variants by predicted phase (in *trans*, unphased, and in *cis*) and the second table counts individuals carrying homozygous variants (**Fig. 6b**). We envision that these data will aid the medical genetics community in interpreting the clinical significance of co-occurring variants in the context of recessive conditions. The data for all genes are also available as a downloadable table within gnomAD v2.

## Discussion

In this work, we have leveraged a large exome sequencing cohort to estimate haplotype frequencies for pairs of rare variants within genes, and demonstrate that these haplotype frequency estimates can be utilized to predict phase of pairs of variants. Overall, we achieve high accuracy across a range of allele frequencies and across genetic ancestries, and demonstrate that our approach is able to distinguish variants that are likely compound heterozygous in a clinical setting. Finally, we freely disseminate our results in an easy-to-use browser for the community.

Our work addresses an important challenge of phasing rare variants using short-read exome sequencing data. While this scenario is very common in medical genetics, other phasing approaches such as phasing-by-transmission or population-based phasing are challenging to apply. We leveraged gnomAD to estimate haplotype frequencies to predict phase of variant pairs seen within genes. We found that our approach was able to address these challenges of phasing in medical genetics and was generally accurate across a range of AFs (even for singleton variants) and across genetic ancestries. Most notably, 96.9% of rare (AF < 5%) variant pairs in a given individual had both variants present in gnomAD and therefore were amenable to phasing using our approach, which is much higher than the proportion amenable to phasing using physical read data. Overall, 92.3% of variant pairs observed in a given individual were correctly phased using our approach. Thus, our approach can be applied to the vast majority of rare variant pairs and can generate accurate phasing estimates for variants of medical importance in rare recessive genetic diseases.

However, we found that our approach was less accurate for rare variants in *cis*. This lower accuracy for variants in *cis* is intuitive, as for rare variants, a recombination event or germline mutation event is much more likely to disrupt a haplotype comprised of two variants than to bring two rare variants onto the same haplotype. Consistent with this intuition, we found that for variants in *cis,* phasing accuracy diminished linearly with genetic distance as a measure of recombination rates, but that phasing accuracy was maintained across genetic and physical distances between pairs of variants in *trans*. Similarly, the phasing accuracy for variant pairs in *cis* was lower at more mutable sites such as CpG sites that are frequently methylated. Thus, users should exercise caution for rare variants at highly mutable sites where our approach predicts the variants to be *trans* since the variants may actually be in *cis*. We note that in the context of recessive disorders in medical genetics, the incorrect phasing of *cis* variants as *trans* is the more desirable error mode to enable follow-up of a variant pair that may be causal.

We also compared population-specific estimates with phasing estimates derived from samples across all genetic ancestry groups in gnomAD v2 (“cosmopolitan”). While population-specific phasing estimates are more likely to match the haplotypes seen in a given individual, they utilize information from fewer samples in gnomAD. We found that, in general, population-specific estimates were similar in accuracy to using cosmopolitan estimates. For individuals of AFR genetic ancestry, however, we found that use of cosmopolitan estimates resulted in slightly lower phasing accuracy than the use of AFR-specific estimates for variants in *trans* (**Fig. 3a)**. This is consistent with the observation that there are more unique haplotypes seen in individuals of AFR genetic ancestry and/or older haplotypes in individuals of AFR genetic ancestry for which recombination is more likely to have occurred^28^. Moreover, there are other genetic ancestry groups not currently represented in gnomAD for which we expect this phasing approach to have lower accuracy than in the well-represented genetic ancestry groups.

Additionally, we recognize that many individuals are not well represented by a discrete genetic ancestry group, but instead represent admixtures of two or more populations. Future work on phasing will likely benefit from considering ancestry as a continuous variable^29^. For the analysis of rare disease patients with candidate compound heterozygous variants, our data suggests that population-specific estimates, when available, should be used first-line followed by the cosmopolitan estimate.

Pairs of singleton variants pose a unique challenge. When a pair of singleton variants is observed in different individuals in a population, this provides evidence that the variants are on different haplotypes. However, if a pair of singleton variants is observed in the same individual in the population, we cannot readily distinguish whether the variants are on the same haplotype or different haplotypes as we lack information from other individuals in the population for singleton variants. For this reason, we have chosen to not report phasing estimates for singleton variant pairs that are observed in the same individuals in gnomAD. Nonetheless, using our trio data, 93% of these singleton variant pairs observed in the same individual in gnomAD were in *cis* based on our trio validation data.

Our work focuses on the challenging, yet common, scenario of determining phase for rare variants identified in exome sequencing of rare disease patients in the absence of parental data. We utilized the EM algorithm to phase pairs of variants instead of more sophisticated population-based phasing approaches for several reasons^16–19^. First, exome and targeted gene panel sequencing data pose a unique challenge for population-based phasing given the sparsity of variants, precluding the use of common non-coding variants as a “scaffold” for phasing.

Recent work performed population-based phasing of rare variants from exome sequencing data by combining the exome data with SNP genotyping arrays^19,30^. However, SNP genotyping data are not usually generated in conjunction with a clinical sequencing test and were not readily available for much of gnomAD. Second, rare variants, which are of the greatest interest in Mendelian diseases, are challenging to phase using population-based approaches given the small numbers of shared haplotypes from which to derive phasing estimates in the population.

Recent methods have shown accurate phasing of rare variants using genome sequencing data^17,19,31^, but relies on a large genome reference panel. In our work, there were limited numbers of genome sequences available for use in a population-based phasing approach. However, as the numbers of genome sequencing samples increases in future releases of gnomAD, this may represent a tractable and more accurate approach for phasing of rare variants. Even with the falling cost of genome sequencing, exome sequencing and targeted gene sequencing remain commonly used in clinical settings. Thus, we anticipate that our approach and the resources we have generated will remain useful. Third, we found that application of the EM algorithm to pairs of variants was more intuitive to illustrate how phasing estimates were derived from genotype data, allowing users to more easily assess the reliability of phasing estimates we provide for any given pair of variants. Together, the EM algorithm provided us with the unique ability to phase pairs of rare variants in exome data in an intuitive fashion.

Utilizing phase estimates to tabulate the number of individuals in gnomAD with two rare predicted in *trans* variants by gene, we found that there are only a small number of “human knock-out” genes affected by predicted compound heterozygous (in *trans*) loss-of-function variants, and that this number is substantially lower than is observed for homozygous loss-of-function variants. These compound heterozygous “human knock-out” events occurred in genes that are not known to be essential, an expected finding given that gnomAD is largely depleted of individuals with severe and early-onset conditions. When analyzing the 23,667 individuals that carry two pLoF variants with AF ≤ 1%, we predict that in 20,706 (88%) of those individuals, the variant pair is in *cis* and only a small fraction (∼0.2%) are confidently predicted to be in *trans*. This may be counter-intuitive, so warrants emphasis: when a pair of rare pLoF variants is observed in the same gene in an individual from a general population sample, it is vastly more likely that these variants are carried on the same haplotype than that the individual is a genuine “knock-out” due to compound heterozygosity.

To aid the medical genetics community in interpreting the clinical significance of rare co-occurring variants in the context of recessive disease, we have released these counts across a spectrum of variant consequences (pLoF, missense, and synonymous) and allele frequencies by gene in the gnomAD browser. These provide background frequencies–to the best of our ability–of compound heterozygous rare damaging variants. These background frequencies can be used to assess the probability that a given variant pair identified in a patient may have occurred by chance. We note, however, that our ability to identify rare variant pairs in *trans* in gnomAD v2 individuals is limited by the fact the same dataset was used for training. Indeed, in these individuals, our ability to detect variant pairs in *trans* extends largely to variant pairs with AF > 0.5%, as nearly all rarer variant combinations were dominated by indeterminate phase and very few predictions for variant pairs in *trans*. The per-gene variant co-occurrence resource developed and released here is therefore to be considered a first step in this space. We plan to use the predictions from this dataset (gnomAD v2) on newer versions of gnomAD with additional samples, where we can more confidently predict rare variant pairs that are in *trans*. Beyond our current restriction to predicting within the gnomAD dataset, there are several other important limitations to our work. First, we have only reported phasing estimates for rare coding and flanking intronic/UTR variant pairs within genes in order to limit the computational burden. We believe that these are the variant pairs of most interest to the medical genetics community, but acknowledge that phase with deeper intronic variation will become important as more genome sequencing is performed. Second, while there was a broad range of genetic ancestral diversity represented in our samples, future studies would benefit from even larger sample sizes, especially for genetic ancestry groups not well represented in our present study. Finally, we have only tested our phasing accuracy in a clinical setting in a retrospective manner and future prospective studies will be needed to confirm the clinical utility of our approach.

## Online Methods

### gnomAD characteristics and data processing

In this work, we used exome sequencing data from the gnomAD v2.1 dataset (n = 125,748 individuals). These data were uniformly processed, underwent joint variant calling, and rigorous quality control, as described in Karczewski et al.^20^. Briefly, we aggregated ∼200k exome sequences and ∼20k genome sequences, primarily from case-control studies of common adult-onset conditions, and applied a BWA-Picard-GATK pipeline^32^. Using Hail (https://github.com/hail-is/hail), we then removed samples that (1) failed population- and platform-specific quality control, (2) had second-degree or closer relations in the dataset, (3) did not have appropriate consent for release, and (4) had known severe, early-onset conditions. For variant quality control, we trained a random forest on site-level and genotype-level metrics (e.g., the quality by depth, QD), and demonstrated that it achieved both high precision and recall for both common and rare variants.

We subsetted the final cleaned gnomAD dataset for variants with global AF in gnomAD exomes < 5% that were either coding, flanking intronic (from position −1 to −3 in acceptor sites, and +1 to +8 in donor sites) or in the 5’/3’ UTRs. In total, this encompasses 5,320,037,963 unique variant pairs across 19,877 genes when removing singleton-singleton variant pairs seen in the same individual. We specifically extracted 20,921,100 pairs of variants, most of which were seen at least once in the same individual to create a more manageable downloadable file.

Analysis in this manuscript was performed using Hail version 0.2.105^33^, and analysis code is available at https://github.com/broadinstitute/gnomad_chets.

### Haplotype estimates

Consider two variants, A and B. A and B represent the major alleles, and a and b represent the respective minor alleles. There are thus 9 pairwise genotypes for A and B: AABB, AaBB, aaBB, AABb, AaBb, aaBb, AAbb, Aabb, and aabb. Of these pairwise genotypes, only the phase for the double heterozygote (AaBb) is unknown. From these 9 possible genotypes, there are four possible haplotype configurations: AB, Ab, aB, and ab.

For each pair of variants, we applied the expectation-maximization (EM) algorithm^21^ to estimate haplotype frequencies from genotype counts. The initial conditions of the EM algorithm were set by partitioning the doubly heterozygous (AaBb) genotype counts equally between the AB|ab and Ab|aB haplotype configurations. The EM algorithm was run until convergence or until the absolute value of the difference between consecutive maximum likelihood function values was less than 1×10^−7^. We calculated haplotype frequencies based on genotypes present within the same genetic ancestry group (“population-specific”) or using all samples from gnomAD (“cosmopolitan”). Haplotype frequency estimates were performed using Hail.

We then calculate P_trans_ as the likelihood that any given pair of doubly heterozygous variants (AaBb) in a patient is compound heterozygous (Ab|aB). P_trans_ can be calculated simply from the haplotype frequency estimates (AB, Ab, aB, and ab):

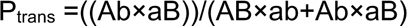

Thus, P_trans_ simply represents the probability that the patient is compound heterozygous by inheriting both the Ab and aB haplotypes.

### Determination of P_trans_ cutoffs

To determine P_trans_ cutoffs for classifying variants as occurring in *cis* or *trans*, we binned variant pairs on odd chromosomes (chromosome 1, 3, 5, etc) in increments of 0.01 of P_trans_. For each bin, we calculated the proportion of variant pairs in that bin that are compound heterozygous based on phasing by trio data. The P_trans_ for variants in *trans* was determined as the minimum P_trans_ such that 90% of variants in the bin are compound heterozygous based on trio data. The P_trans_ for variants in *cis* was determined as the maximum P_trans_ such that 90% of variants in the bin are in *cis* based on trio data. For these calculations, we used only variants where both variants had a population AF ≥ 1×10^−4^. We used trio samples across all genetic ancestry groups and population-specific P_trans_ values for determination of the cutoffs.

### Trio validation data

We made use of 4,992 parent-child trios that were jointly processed and variant-called with gnomAD. Having access to parental genotypes allows us to perform phase-by-transmission and accurately determine whether two co-occurring variants in the same gene are in *cis* or in *trans*. First, we estimated genetic ancestry of each individual in the trios by using ancestry inference estimates from the full gnomAD dataset, as previously described^20^. Briefly, we selected bi-allelic variants that passed all hard filters, had allele frequencies in a joint exome and genome callset > 0.001, and high joint call rates (> 0.99). The variants were then LD-pruned (r^2^ = 0.1) and used in a principal component analysis (PCA). We previously used samples with known genetic ancestry to train a random forest on the first 20 principal components (PCs), and assigned samples to a genetic ancestry group based on having a random forest probability > 0.9. For the trios in this cohort, we projected their PCs for genetic ancestry onto the same gnomAD v2 samples to infer the genetic ancestry used here (**Figure S1**). Of these 4922 trios, 4,775 of the children from the trios were assigned to one of the seven genetic ancestry groups in this study based on PCA and were used in this study.

We then phased the trio data using the Hail *phase_by_transmission* (https://hail.is/docs/0.2/experimental/index.html#hail.experimental.phase_by_transmission) function, which uses Mendelian transmission of alleles to infer haplotypes in trios for all sites that are not heterozygous in all members of the trio. Assigning haplotypes in trios based on parental genotype has traditionally been the gold standard, has switch error rates below 0.1%, and importantly errors aren’t dependent on the allele frequency of the variants phased^34^. To maximize our confidence in the genotypes and phasing, we filtered genotypes to include only those with genotype quality (GQ) > 20, depth > 10 and allele balance > 0.2 for heterozygous genotypes prior to phasing. Sex chromosomes were excluded. In total, there were 339,857 unique variant pairs and 1,115,347 total variant pairs.

We compared trio phasing-by-transmission with phasing using gnomAD on even chromosomes (e.g., chromosomes 2, 4, 6, etc). Of these 4,775 trio samples, 3,836 were in the full release of gnomAD and were removed from gnomAD for trio validation. This resulted in a set of 121,912 gnomAD samples from which we derived haplotype estimates. We then performed phasing using the EM algorithm as above.

We classified trio variant pairs into 1) unable to phase using our approach (either variant not seen in gnomAD, or singleton-singleton variant pairs seen in the same individual in gnomAD), 2) indeterminate phase (those with intermediate 0.02 < P_trans_ < 0.55), 3) incorrectly phased, or 4) correctly phased. Accuracy was calculated as the number of variant pairs correctly phased divided by the number of pairs correctly and incorrectly phased.

### CpG analysis

Single nucleotide variants seen in the trio data were divided into transitions and transversions. Transitions were further subdivided into those that are CpG mutations (5’-CpG-3’ mutating to 5’-TpG-3’) and those that are not. For each CpG transition, we calculated the mean DNA methylation values across 37 tissues in ENCODE^20^. We then stratified CpG transitions into 3 levels: low (missing or < 0.2), medium (0.2–0.6), and high (> 0.6) methylation. Phasing accuracy–here, the proportion correct (correct classifications/all classifications)–was then calculated for pairwise combinations of transversions, non-CpG transitions, low methylation CpG transitions, medium methylation CpG transitions, and high methylation CpG transitions. All SNVs were included in the analysis and population-specific EM estimates were used.

### Calculating accuracy as a function of genetic distance

To estimate the genetic distance between pairs of genetic variants, we interpolated genetic distances between variant pairs using a genetic map from HapMap2^35^ (https://github.com/joepickrell/1000-genomes-genetic-maps). We downloaded a pre-generated HapMap2 genetic map representing average over recombination rates in the CEU, YRI, and ASN populations. We then ran interpolate_maps.py (downloaded from https://github.com/joepickrell/1000-genomes-genetic-maps/blob/master/scripts/interpolate_maps.py) for all variant pairs in the phasing trio data. As above, accuracy is the proportion of correct classifications.

### MNV analysis

We obtained multi-nucleotide variant pairs for which read-back phasing had previously been calculated^22^. We included only multi-nucleotide variant pairs where each constituent variant was analyzed in our study. Phasing estimates were calculated using cosmopolitan EM estimates.

### Rare disease patient analysis

627 patients from the Broad Institute Center for Mendelian Genetics (CMG)^25^ with a confident or strong candidate genetic diagnosis of a Mendelian condition were selected for analysis. Each patient carried two presumed bi-allelic variants in an autosomal recessive disease gene consistent with the patient’s phenotype. For 293 of the patients, both variants were present in gnomAD and phase was predicted. Trio-sequencing (i.e., sequencing of the proband and the two unaffected biological parents) had been performed for 168 of the 293 patients. For fully sequenced trios, we were able to confirm phasing of the two variants via phase-by-transmission.

### Determining counts of individuals with two rare, damaging variants

Variants were annotated with the worst consequence on the canonical transcript by the Ensembl Variant Effect Predictor (VEP)^36^. pLoF were annotated with LOFTEE^20^, and only high confidence LoF variants were counted as “pLoF”. Missense variants were annotated with REVEL^26^. REVEL scores ≥ 0.932 were counted as “strong_revel_missense”, ≥ 0.773 as “moderate_to_strong_revel_missense”, ≥ 0.644 as “weak_to_strong_revel_missense” in line with recent ClinGen recommendations^27^.

Variant pairs were annotated with predicted phase based on the P_trans_ thresholds. All singleton-singleton variant pairs (AC = 1) and variant pairs with an indeterminate P_trans_ values (0.02 < P_trans_ < 0.55) were annotated as unphased.

Five AF thresholds were selected for analysis and variant pairs were filtered based on the highest global AF and, where available, the “popmax” AF of each variant in gnomAD (i.e., the highest AF information for the non-bottlenecked population - excluding ASJ, FIN and “Remaining”): 0.5%, 1%, 1.5%, 2%, and 5%. Further, all variant pairs containing a variant with an AF > 5% in a bottlenecked population were filtered out.

The number of individuals carrying a variant pair (irrespective of phase) and the number indicated to be compound heterozygous (in *trans*), unphased (indeterminate), and on the same haplogroup (in *cis*) were counted gene-wise by AF threshold and combined functional consequences (26 consequences). This counting was repeated twice, once restricting individuals to be counted in only one phase group, prioritizing in *trans* over unphased and unphased over in *cis* (displayed in the “variant co-occurrence” gnomAD browser feature), and once allowing individuals to be counted in multiple phase groups, if carrying multiple variant pairs in the same gene with different phase predictions.

### Essential gene lists

The following essential gene lists were queried for the presence of the true “human knock-out” genes identified in this study:

● 2,454 genes essential in mice from Georgi et al. 2013^37^
● 553 pan-cancer core fitness genes from Behan et al., 2019^38^
● 360 core essential genes from genomic perturbation screens from Hart et al. 2014^39^
● 684 genes essential in culture by CRISPR screening from Hart et al. 2017^40^
● 1,075 genes annotated by the ADaM analysis of a large collection of gene dependency profiles (CRISPR-Cas9 screens) across human cancer cell lines from Vinceti et al. 2021^41^

## Code availability

The code used to estimate P_trans_ estimates for variant pairs and to determine the number of individuals carrying rare, compound heterozygous variants can be found at: https://github.com/broadinstitute/gnomad_chets

## Data availability

We provide both web-based look up tools and downloads for the data generated here. A look-up tool to find the likely co-occurrence pattern between two rare (global allele frequency in gnomAD exomes < 5%) coding, flanking intronic (from position −1 to −3 in acceptor sites, and +1 to +8 in donor sites) or 5’/3’ UTR variants can be found at: https://gnomad.broadinstitute.org/variant-cooccurrence

Additionally, we display the per-gene counts tables that detail the number of individuals with two rare variants, stratified by AF and functional consequence, on each gene’s main page. One table details counts of individuals with two heterozygous variants and includes predicted phase, while the second details individuals with homozygous variants. Both can be found by clicking on the “Variant Co-occurrence” tab on each gene’s main page.

All variant co-occurrence tables can be downloaded from: https://gnomad.broadinstitute.org/downloads#v2-variant-cooccurrence

## Ethical compliance and informed consent statement

Informed consent was obtained by collaborators for all participants in the Broad Institute Center for Mendelian Genetics (CMG), and individual-level data, including genomics and clinical data, were de-identified and coded by our collaborators before analysis here. We have complied with all relevant ethical regulations. This work was approved by the Broad Institute of MIT and Harvard, Mass General Brigham IRB.

## Author contributions

M.H.G., L.C.F., S.L.S., J.K.G., A.O.D.L., K.J.K., D.G.M., and K.E.S. conceived and designed experiments. M.H.G., L.C.F., S.L.S., and J.K.G. performed the analyses. N.A.W., P.W.D., and M.S. developed visualizations for the web browser. E.G. and M.S-B. performed variant curation. G.T., B.M.N., J.N.H., H.L.R., M.J.D., A.O.D.L., and K.J.K. provided data and analysis suggestions. J.N.H., D.G.M., and K.E.S. supervised the work. M.H.G., L.C.F., S.L.S., J.K.G., and K.E.S. completed the primary writing of the manuscript with input and approval of the final version from all other authors.

## Acknowledgements

We would like to thank all members of the gnomAD team for helpful comments and suggestions, and to particularly recognize the members of the gnomAD methods and browser teams who worked hard over many years to provide cleaned datasets, easy-to-use browsers, and visualizations. This work was supported by NHGRI U24HG011450, UM1HG008900, and U01HG011755.

References

## Supplementary Figures

**Supplementary Figure 1:**
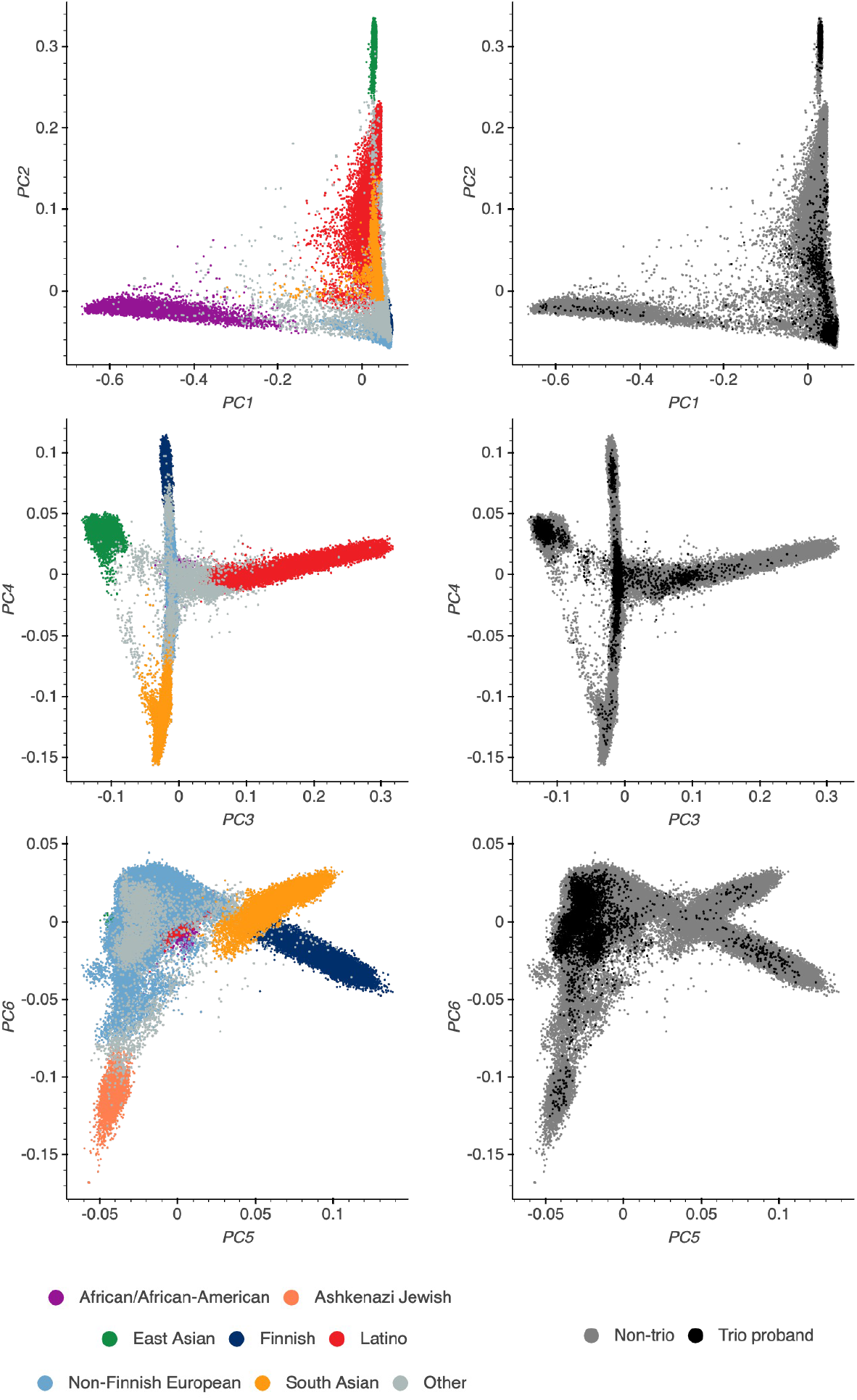
Principal comReferencesponent analysis (PCA) plot for the full gnomAD v2 cohort (left) and specifically for the trios (right, trios in black) included in this paper. The top row shows PC1 vs PC2, the middle row shows PC3 vs PC4, and the bottom row shows PC5 vs PC6. Genetic ancestry group labels for the global gnomAD populations were done as described in Karczewski et al. 2020^20^.

**Supplementary Figure 2:**
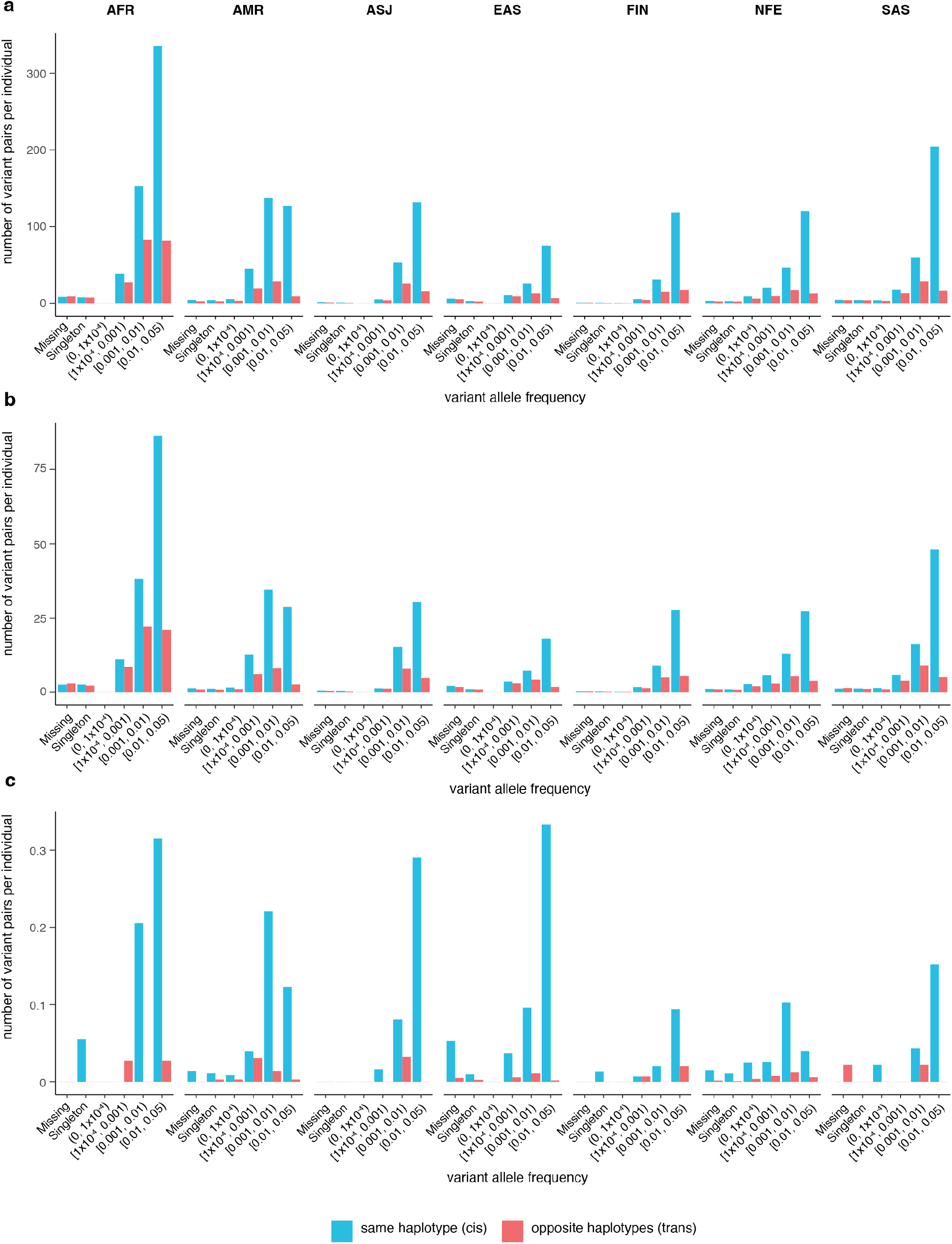
Number of variant pairs observed per trio sample as a function of ancestry and AF. All variant pairs are shown in **a.** Variant pairs in which both variants are moderate effect or predicted loss-of-function (pLoF) are shown in **b.** Variant pairs in which both variants are pLoF are shown in **c.** Variant AF is the AF of the less common variant in a given variant pair and is population-specific frequency. AFR = African/African American; AMR = Admixed American/Latino; ASJ = Ashkenazi Jewish; EAS = East Asian; FIN = Finnish; NFE = non-Finnish European; SAS = South Asian.

**Supplementary Figure 3:**
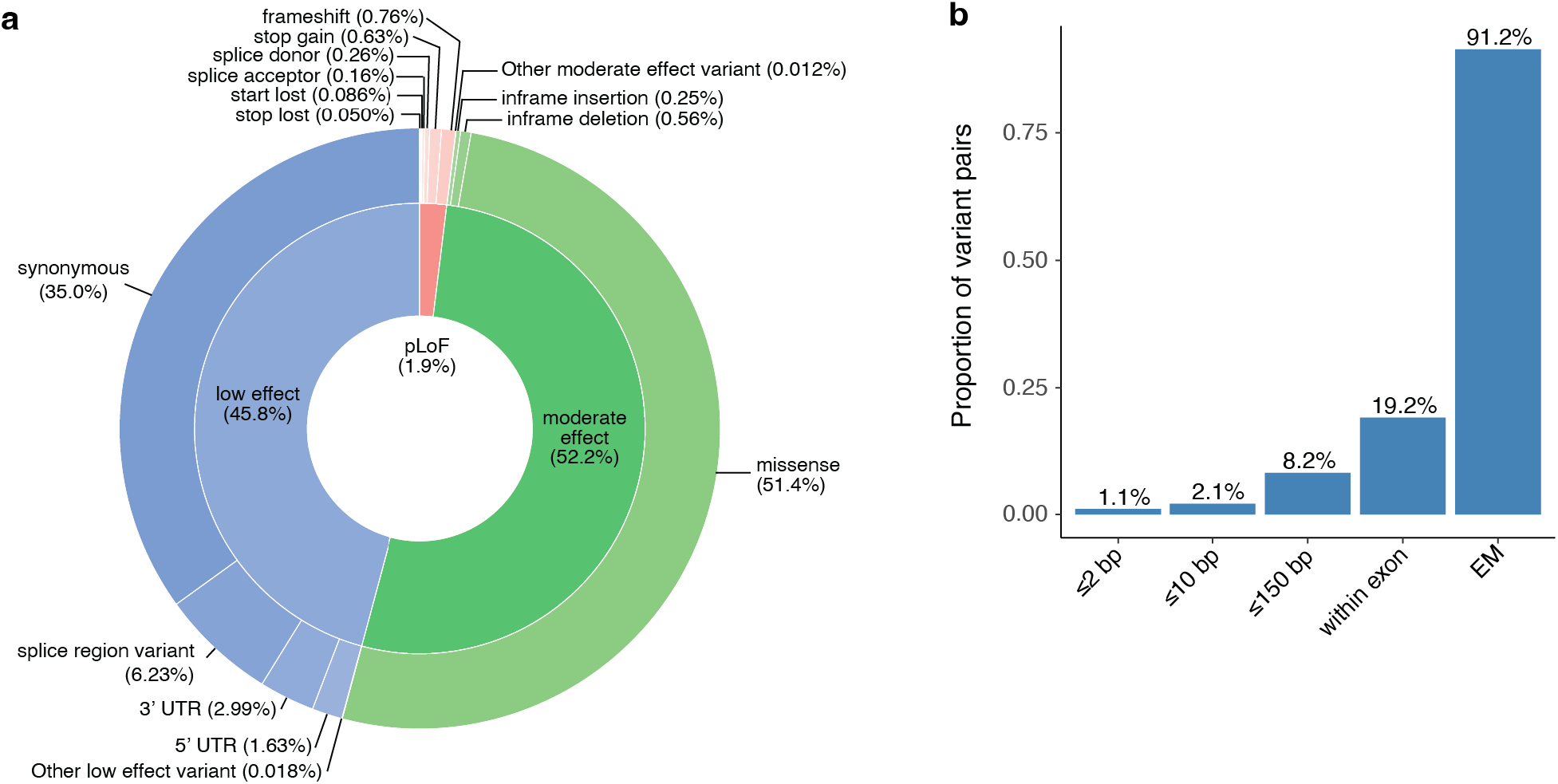
**a,** Pie chart of variant effect annotations in the trio samples. Effect predictions are stratified among pLoF, moderate effect, and low effect variants. Percentages are shown in parentheses. **b,** Proportion of variant pairs falling within 2 bp, within 10 bp, within 150 bp, within the same exon, and proportion that can be phased using the EM algorithm and the gnomAD resource.

**Supplementary Figure 4:**
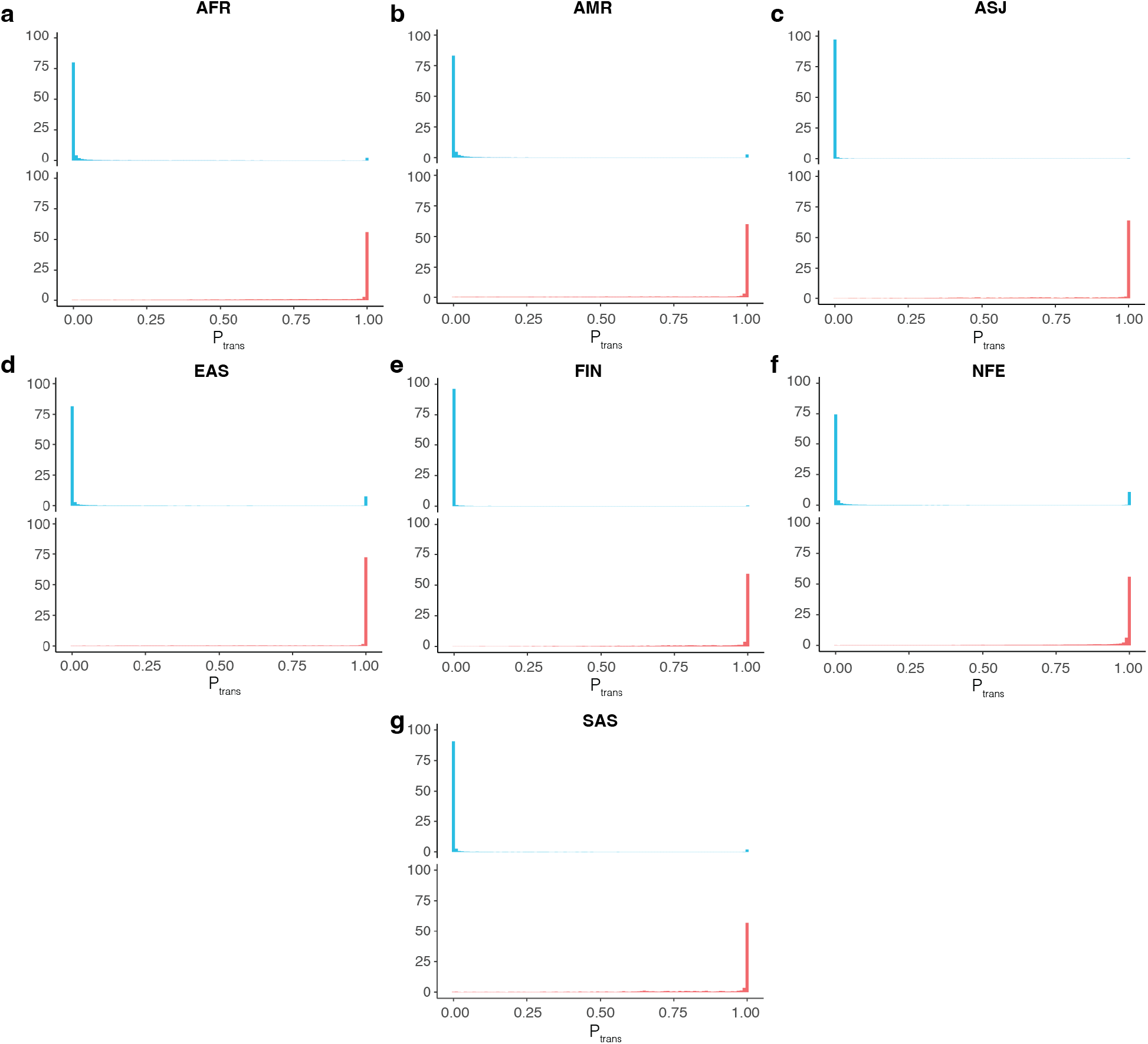
**a-g,** Histogram of P_trans_ scores for variant pairs in *cis* (top, blue) and in *trans* (bottom, red) for each population. P_trans_ scores are population-specific. AFR = African/African American; AMR = Admixed American/Latino; ASJ = Ashkenazi Jewish; EAS = East Asian; FIN = Finnish; NFE = non-Finnish European; SAS = South Asian.

**Supplementary Figure 5:**
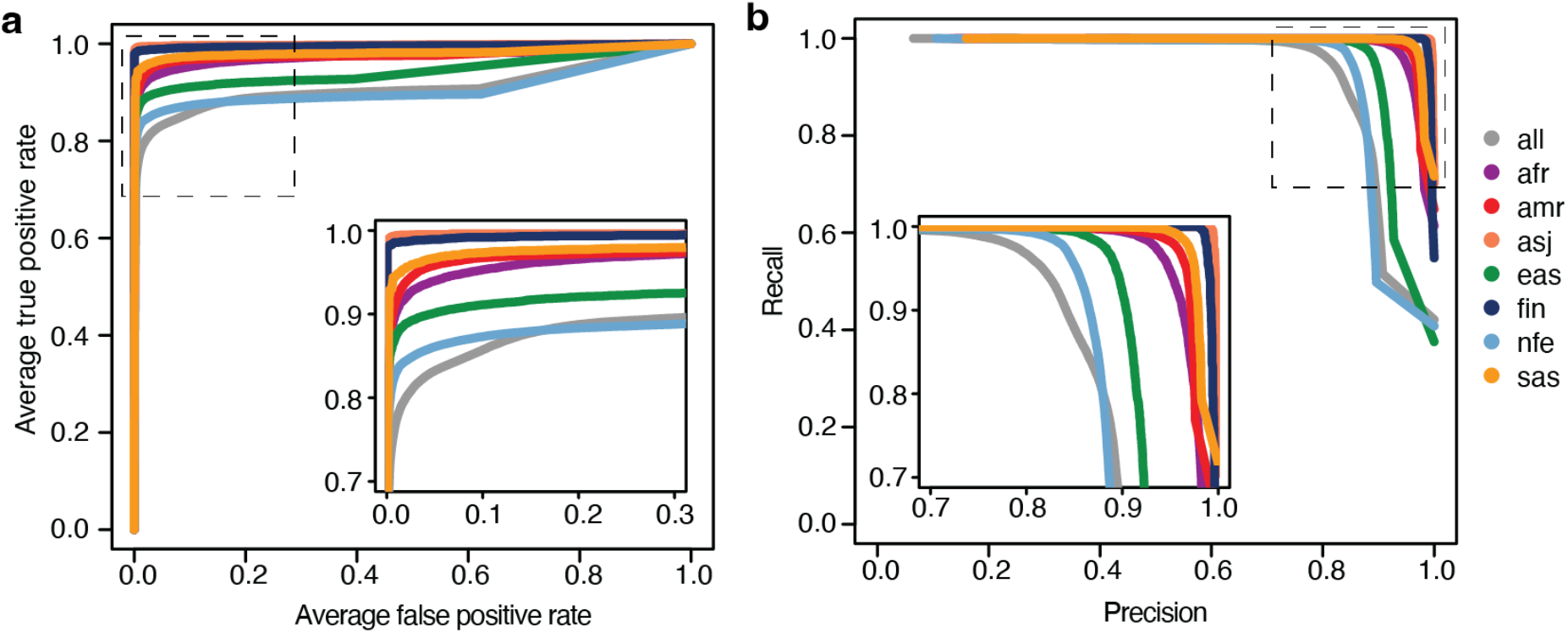
Receiver-operator (**a**) and Precision-recall (**b**) curves for use of P_trans_ for distinguishing between variant pairs on same versus opposite haplotypes. Separate lines are shown for each genetic ancestry group. P_trans_ scores are population-specific. AFR = African/African American; AMR = Admixed American/Latino; ASJ = Ashkenazi Jewish; EAS = East Asian; FIN = Finnish; NFE = non-Finnish European; SAS = South Asian.

**Supplementary Figure 6:**
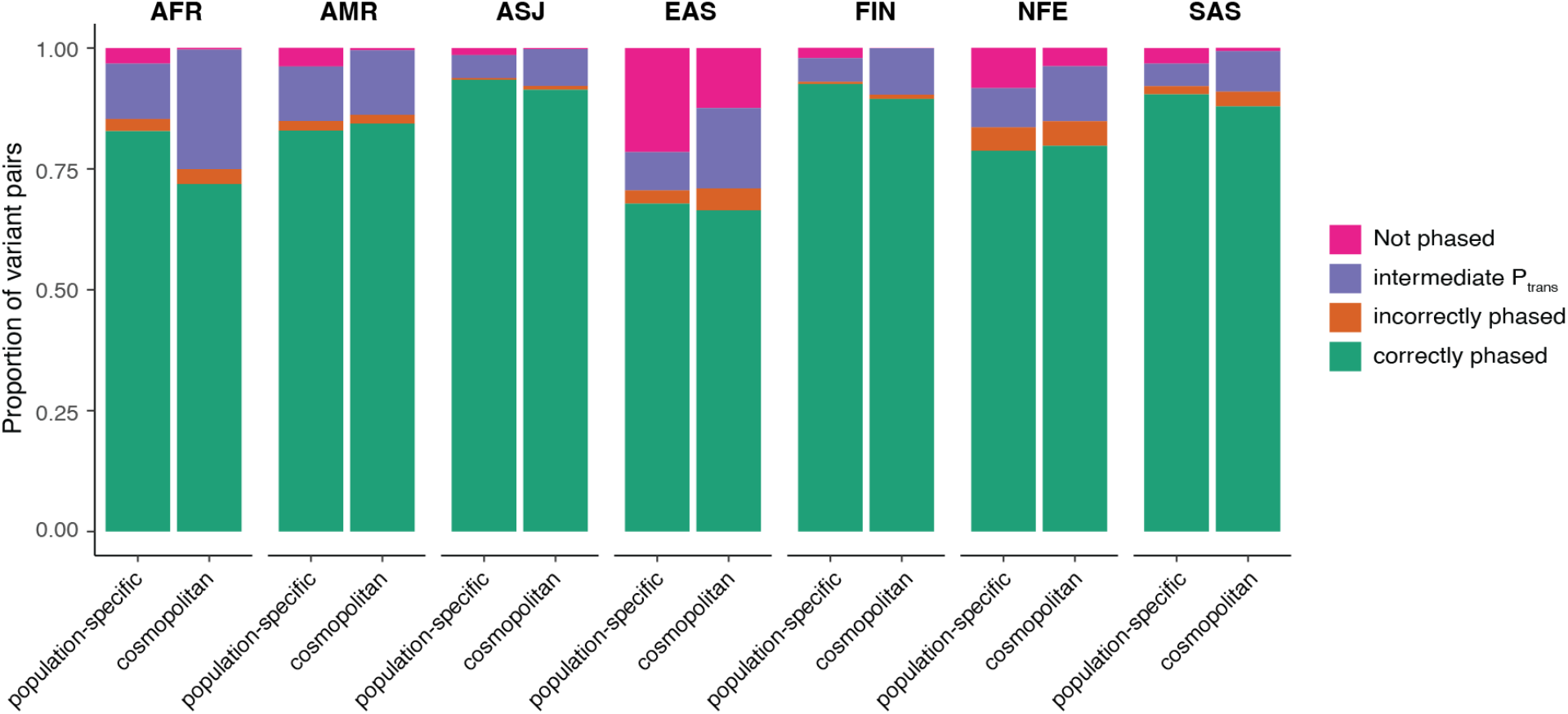
Phasing performance for population-specific versus cosmopolitan P_trans_ scores for each population. AFR = African/African American; AMR = Admixed American/Latino; ASJ = Ashkenazi Jewish; EAS = East Asian; FIN = Finnish; NFE = non-Finnish European; SAS = South Asian.

**Supplementary Figure 7:**
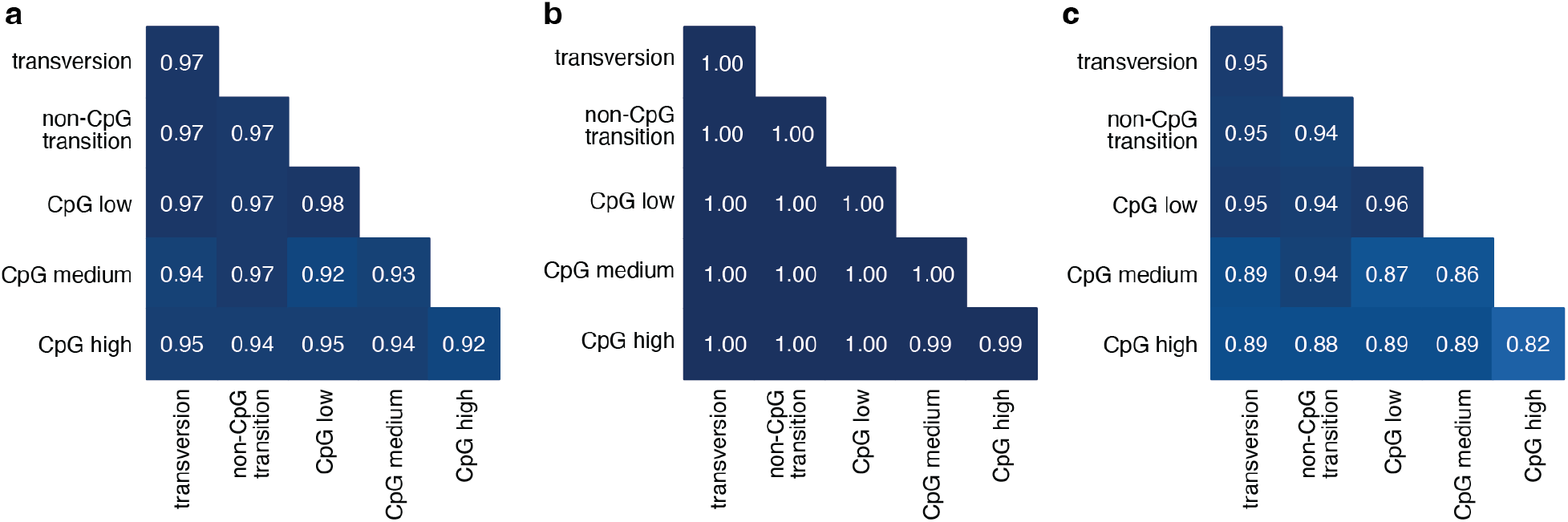
Phasing accuracy for transversions, non-CpG transitions, and CpG transitions. CpG transitions are further stratified by degree of DNA methylation (low, medium, or high) as in Karczewski et al^20^. Shading of squares and numbers in each square represents phasing accuracy. Phasing accuracies are based on variant pairs seen in all populations and utilize population-specific P_trans_ estimates. Accuracy is shown for all variants **(a)**, variants in *trans* **(b),** and variants in *cis* **(c).**

**Supplementary Figure 8:**
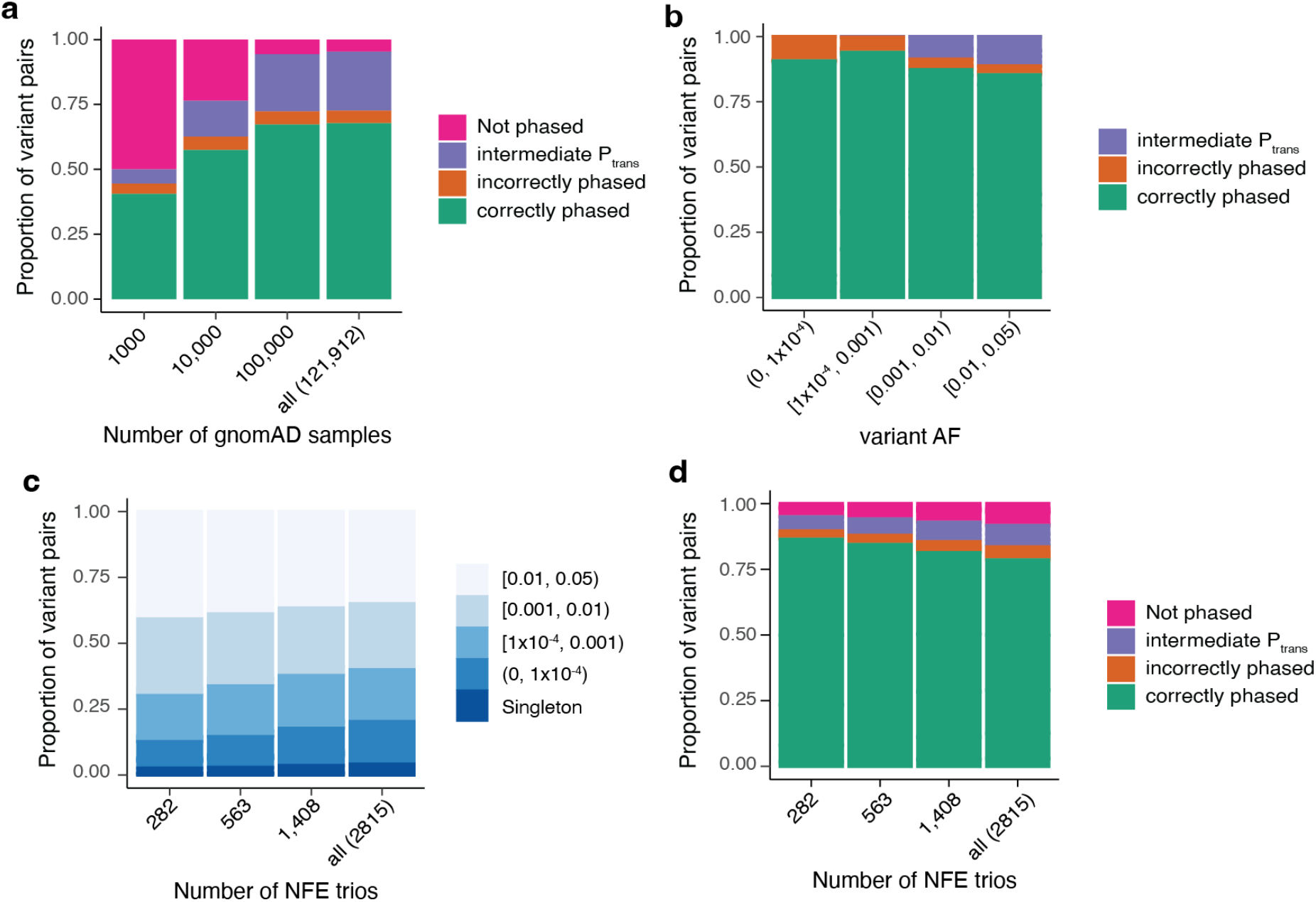
**a,** Phasing performance when subsampling gnomAD to 1000, 10,000, 100,000 or using all samples. Phasing performance is based on cosmopolitan P_trans_ estimates and is calculated across trio samples from all populations. **b,** Phasing performance as a function of variant AF for the more common variant in a variant pair. Phasing performance is based on population-specific P_trans_ estimates and is calculated across trio samples from all populations. **c,** Proportion of variants falling into different AF bins when subsampling NFE gnomAD trios from 2815 trios down to 282, 563, or 1408 trios. Allele frequencies reflect the rarer variant in a variant pair. **d,** Phasing performance when subsampling NFE gnomAD samples as described in **c.**

**Supplementary Figure 9:**
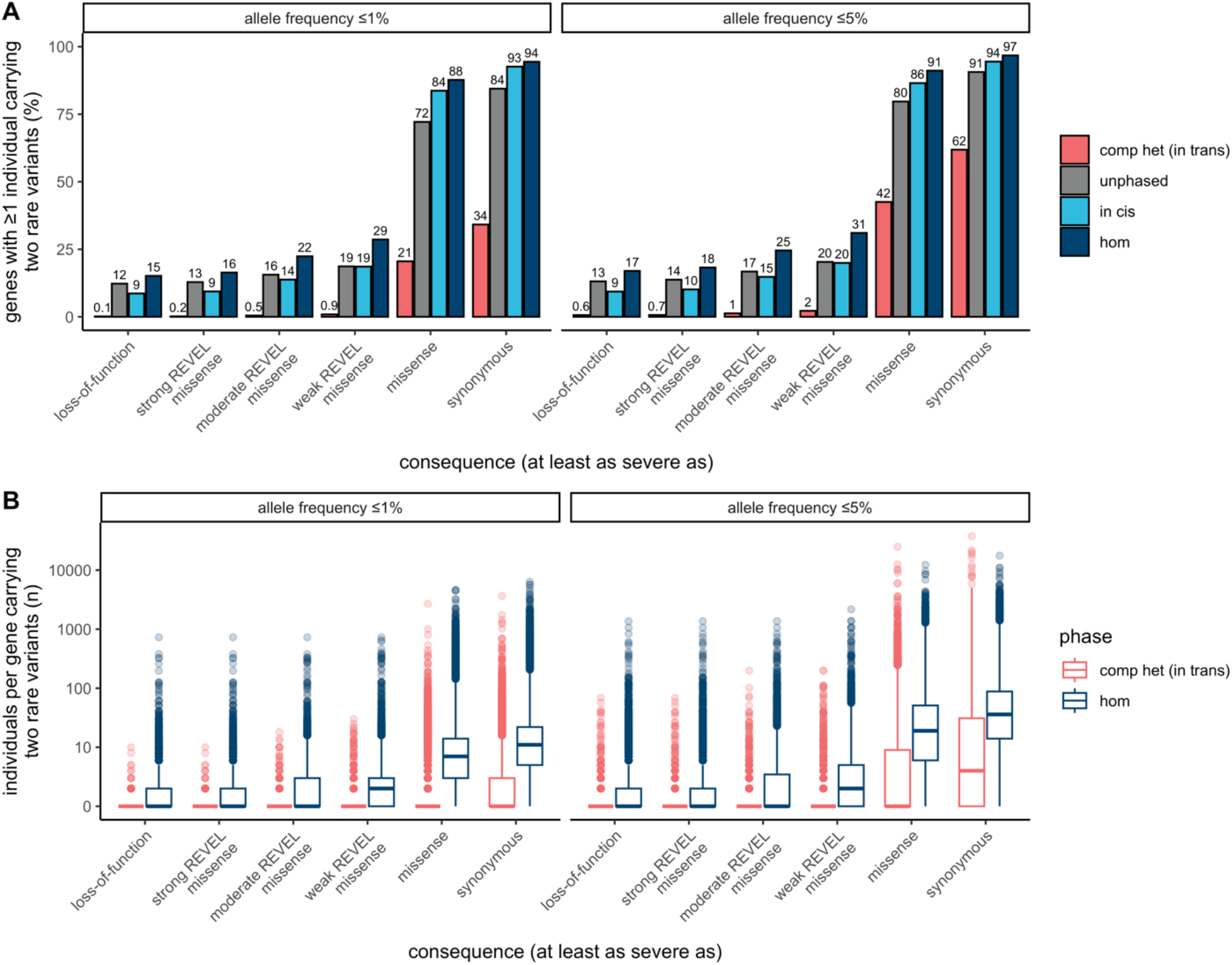
**a,** Proportion of genes with one or more individuals in gnomAD carrying two rare variants at ≤ 1% and ≤ 5% AF stratified by predicted functional effect and phase. For compound heterozygous (comp het, in *trans*), unphased, and in *cis*, both variants in the variant pair must be annotated with a consequence at least as severe as the consequence displayed. **b,** Number of individuals per gene in gnomAD carrying two rare variants at ≤ 1% and ≤ 5% AF stratified by predicted functional effect and phase. For compound heterozygous (in *trans*) both variants in the variant pair must be annotated with a consequence at least as severe as the consequence displayed. In the box plots, the center line is the median, the box limits are the upper and lower quartiles, and the whiskers extend to the 1.5x the interquartile range. Any points shown are outliers. “comp het (in trans)” refers to compound heterozygous; “hom” refers to homozygous.

## Supplemental Tables

**Table S1.** CMG diagnostic variants. In this table, we provide details about the presumed bi-allelic causal variants from 293 individuals from the Broad Institute Center for Mendelian Genetics. For each variant pair, we provide the gene symbol (“gene_name”), information about the position and alleles of both variants, whether both of the variants were singletons in gnomAD (“singleton_singleton”) and seen in the same individual or not, the estimated cosmopolitan P_trans_ value, the predicted phase based on the cosmopolitan P_trans_ value (“cosmopolitan_phase_prediction”), the imputed population ancestry of the CMG individual (“imputed_population_ancestry”), the predicted phased based on the population-specific P_trans_ value (“population_specific_phase_prediction”), the known phase from phase by transmission when trio data were available (“phase_by_transmission”), and an explanation for incorrect predictions where applicable (“incorrect_prediction_explanation”).

**Table S2.** Manual curation results for compound heterozygous loss-of-function variants. Here, we provide the variant curation information for the 28 genes that have predicted compound heterozygous loss-of-function variants with AF ≤ 1%. For every predicted compound heterozygous variant pair, we provide the gene symbol, maximum AF in the gnomAD exomes from the two variants (“variant_pair_max_af”), the number of individuals who carry the variant pair (“n_individuals”), information about the position and alleles of both variants, any manual curation flags e.g., mapping error for the variants, and the final loss-of-function curation for both variants as well as the variant pair (“high_confidence_human_knock_out”).

**Genome Aggregation Database Consortium**

**Maria Abreu^1^, Carlos A. Aguilar Salinas^2^, Tariq Ahmad^3^, Christine M. Albert^4,5^, Jessica Alföldi^6,7^, Diego Ardissino^8^, Irina M. Armean^6,7,9^, Gil Atzmon^10,11^, Eric Banks^12^, John Barnard^13^, Samantha M. Baxter^6^, Laurent Beaugerie^14^, Emelia J. Benjamin^15,16,17^, David Benjamin^12^, Louis Bergelson^12^, Michael Boehnke^18^, Lori L. Bonnycastle^19^, Erwin P. Bottinger^20^, Donald W. Bowden^21,22,23^, Matthew J. Bown^24,25^, Steven Brant^26^, Sarah E. Calvo^6,27^, Hannia Campos^28,29^, John C. Chambers^30,31,32^, Juliana C. Chan^33^, Katherine R. Chao^6^, Sinéad Chapman^6,7,34^, Daniel Chasman^4,35^, Siwei Chen^6,7^, Rex L. Chisholm^36^, Judy Cho^20^, Rajiv Chowdhury^37^, Mina K. Chung^38^, Wendy K. Chung^39,40,41^, Kristian Cibulskis^12^, Bruce Cohen^35,42^, Ryan L. Collins^6,27,43^, Kristen M. Connolly^44^, Adolfo Correa^45^, Miguel Covarrubias^12^, Beryl Cummings^6,43^, Dana Dabelea^46^, Mark J. Daly^6,7,47^, John Danesh^37^, Dawood Darbar^48^, Joshua Denny^49^, Stacey Donnelly^6^, Ravindranath Duggirala^50^, Josée Dupuis^51,52^, Patrick T. Ellinor^6,53^, Roberto Elosua^54,55,56^, James Emery^12^, Eleina England^6^, Jeanette Erdmann^57,58,59^, Tõnu Esko^6,60^, Emily Evangelista^6^, Yossi Farjoun^12^, Diane Fatkin^61,62,63^, Steven Ferriera^64^, Jose Florez^35,65,66^, Laurent C. Francioli^6,7^, Andre Franke^67^, Martti Färkkilä^68^, Stacey Gabriel^64^, Kiran Garimella^12^, Laura D. Gauthier^12^, Jeff Gentry^12^, Gad Getz^35,69,70^, David C. Glahn^71,72^, Benjamin Glaser^73^, Stephen J. Glatt^74^, David Goldstein^75,76^, Clicerio Gonzalez^77^, Julia K. Goodrich^6^, Leif Groop^78,79^, Sanna Gudmundsson^6,7,80^, Namrata Gupta^6,64^, Andrea Haessly^12^, Christopher Haiman^81^, Ira Hall^82^, Craig Hanis^83^, Matthew Harms^84,85^, Mikko Hiltunen^86^, Matti M. Holi^87^, Christina M. Hultman^88,89^, Chaim Jalas^90^, Thibault Jeandet^12^, Mikko Kallela^91^, Diane Kaplan^12^, Jaakko Kaprio^79^, Konrad J. Karczewski^6,7,34^, Sekar Kathiresan^27,35,92^, Eimear Kenny^89,93^, Bong-Jo Kim^94^, Young Jin Kim^94^, George Kirov^95^, Zan Koenig^6^, Jaspal Kooner^31,96,97^, Seppo Koskinen^98^, Harlan M. Krumholz^99^, Subra Kugathasan^100^, Soo Heon Kwak^101^, Markku Laakso^102,103^, Nicole Lake^104^, Trevyn Langsford^12^, Kristen M. Laricchia^6,7^, Terho Lehtimäki^105^, Monkol Lek^104^, Emily Lipscomb^6^, Christopher Llanwarne^12^, Ruth J.F. Loos^20,106^, Steven A. Lubitz^6,53^, Teresa Tusie Luna^107,108^, Ronald C.W. Ma^33,109,110^, Daniel G. MacArthur^6,111,112^, Gregory M. Marcus^113^, Jaume Marrugat^55,114^, Alicia R. Martin^6^, Kari M. Mattila^105^, Steven McCarroll^34,115^, Mark I. McCarthy^116,117,118^, Jacob McCauley^119,120^, Dermot McGovern^121^, Ruth McPherson^122^, James B. Meigs^6,35,123^, Olle Melander^124^, Andres Metspalu^125^, Deborah Meyers^126^, Eric V. Minikel^6^, Braxton D. Mitchell^127^, Vamsi K. Mootha^6,128^, Ruchi Munshi^12^, Aliya Naheed^129^, Saman Nazarian^130,131^, Benjamin M. Neale^6,7^, Peter M. Nilsson^132^, Sam Novod^12^, Anne H. O’Donnell-Luria^6,7,80^, Michael C. O’Donovan^95^, Yukinori Okada^133,134,135^, Dost Ongur^35,42^, Lorena Orozco^136^, Michael J. Owen^95^, Colin Palmer^137^, Nicholette D. Palmer^138^, Aarno Palotie^7,34,79^, Kyong Soo Park^101,139^, Carlos Pato^140^, Nikelle Petrillo^12^, William Phu^6,80^, Timothy Poterba^6,7,34^, Ann E. Pulver^141^, Dan Rader^130,142^, Nazneen Rahman^143^, Heidi L. Rehm^6,27^, Alex Reiner^144,145^, Anne M. Remes^146^, Dan Rhodes^6^, Stephen Rich^147,148^, John D. Rioux^149,150^, Samuli Ripatti^79,151,152^, David Roazen^12^, Dan M. Roden^153,154^, Jerome I. Rotter^155^, Valentin Ruano-Rubio^12^, Nareh Sahakian^12^, Danish Saleheen^156,157,158^, Veikko Salomaa^159^, Andrea Saltzman^6^, Nilesh J. Samani^24,25^, Kaitlin E. Samocha^6,7,27^, Jeremiah Scharf^6,27,34^, Molly Schleicher^6^, Heribert Schunkert^160,161^, Sebastian Schönherr^162^, Eleanor Seaby^6^, Cotton Seed^7,34^, Svati H. Shah^163^, Megan Shand^12^, Moore B. Shoemaker^164^, Tai Shyong^165,166^, Edwin K. Silverman^167,168^, Moriel Singer-Berk^6^, Pamela Sklar^169,170,171^, J. Gustav Smith^152,172,173^, Jonathan T. Smith^12^, Hilkka Soininen^174^, Harry Sokol^175,176,177^, Matthew Solomonson^6,7^, Rachel G. Son^6^, Jose Soto^12^, Tim Spector^178^, Christine Stevens^6,7,34^, Nathan Stitziel^82,179^, Patrick F. Sullivan^88,180^, Jaana Suvisaari^159^, E. Shyong Tai^181,182,183^, Michael E. Talkowski^6,27,34^, Yekaterina Tarasova^6^, Kent D. Taylor^155^, Yik Ying Teo^181,184,185^, Grace Tiao^6,7^, Kathleen Tibbetts^12^, Charlotte Tolonen^12^, Ming Tsuang^186,187^, Tiinamaija Tuomi^79,188,189^, Dan Turner^190^, Teresa Tusie-Luna^191,192^, Erkki Vartiainen^193^, Marquis Vawter^194^, Christopher Vittal^6,7^, Gordon Wade^12^, Arcturus Wang^6,7,34^, Qingbo Wang^6,133^, James S. Ware^6,195,196^, Hugh Watkins^197^, Nicholas A. Watts^6,7^, Rinse K. Weersma^198^, Ben Weisburd^12^, Maija Wessman^79,199^, Nicola Whiffin^6,200,201^, Michael W. Wilson^6,7^, James G. Wilson^202^, Ramnik J. Xavier^203,204^, Mary T. Yohannes^6^**

^1^University of Miami Miller School of Medicine, Gastroenterology, Miami, USA

^2^Unidad de Investigacion de Enfermedades Metabolicas, Instituto Nacional de Ciencias Medicas y Nutricion, Mexico City, Mexico

^3^Peninsula College of Medicine and Dentistry, Exeter, UK

^4^Division of Preventive Medicine, Brigham and Women’s Hospital, Boston, MA, USA

^5^Division of Cardiovascular Medicine, Brigham and Women’s Hospital and Harvard Medical School, Boston, MA, USA

^6^Program in Medical and Population Genetics, Broad Institute of MIT and Harvard, Cambridge, MA, USA

^7^Analytic and Translational Genetics Unit, Massachusetts General Hospital, Boston, MA, USA

^8^Department of Cardiology University Hospital, Parma, Italy

^9^European Molecular Biology Laboratory, European Bioinformatics Institute, Wellcome Genome Campus, Hinxton, Cambridge, UK

^10^Department of Biology Faculty of Natural Sciences, University of Haifa, Haifa, Israel

^11^Departments of Medicine and Genetics, Albert Einstein College of Medicine, Bronx, NY, USA

^12^Data Science Platform, Broad Institute of MIT and Harvard, Cambridge, MA, USA

^13^Department of Quantitative Health Sciences, Lerner Research Institute Cleveland Clinic, Cleveland, OH, USA

^14^Sorbonne Université, APHP, Gastroenterology Department Saint Antoine Hospital, Paris, France

^15^NHLBI and Boston University’s Framingham Heart Study, Framingham, MA, USA

^16^Department of Medicine, Boston University Chobanian and Avedisian School of Medicine, Boston, MA, USA

^17^Department of Epidemiology, Boston University School of Public Health, Boston, MA, USA

^18^Department of Biostatistics and Center for Statistical Genetics, University of Michigan, Ann Arbor, MI, USA

^19^National Human Genome Research Institute, National Institutes of Health Bethesda, MD, USA

^20^The Charles Bronfman Institute for Personalized Medicine, Icahn School of Medicine at Mount Sinai, New York, NY, USA

^21^Department of Biochemistry, Wake Forest School of Medicine, Winston-Salem, NC, USA

^22^Center for Genomics and Personalized Medicine Research, Wake Forest School of Medicine, Winston-Salem, NC, USA

^23^Center for Diabetes Research, Wake Forest School of Medicine, Winston-Salem, NC, USA

^24^Department of Cardiovascular Sciences, University of Leicester, Leicester, UK

^25^NIHR Leicester Biomedical Research Centre, Glenfield Hospital, Leicester, UK

^26^John Hopkins Bloomberg School of Public Health, Baltimore, MD, USA

^27^Center for Genomic Medicine, Massachusetts General Hospital, Boston, MA, USA

^28^Harvard School of Public Health, Boston, MA, USA

^29^Central American Population Center, San Pedro, Costa Rica

^30^Department of Epidemiology and Biostatistics, Imperial College London, London, UK

^31^Department of Cardiology, Ealing Hospital, NHS Trust, Southall, UK

^32^Imperial College, Healthcare NHS Trust Imperial College London, London, UK

^33^Department of Medicine and Therapeutics, The Chinese University of Hong Kong, Hong Kong, China

^34^Stanley Center for Psychiatric Research, Broad Institute of MIT and Harvard, Cambridge, MA, USA

^35^Department of Medicine, Harvard Medical School, Boston, MA, USA

^36^Northwestern University Feinberg School of Medicine, Chicago, IL, USA

^37^University of Cambridge, Cambridge, England

^38^Departments of Cardiovascular, Medicine Cellular and Molecular Medicine Molecular Cardiology, Quantitative Health Sciences, Cleveland Clinic, Cleveland, OH, USA

^39^Department of Pediatrics, Columbia University Irving Medical Center, New York, NY, USA

^40^Herbert Irving Comprehensive Cancer Center, Columbia University Medical Center, New York, NY, USA

^41^Department of Medicine, Columbia University Medical Center, New York, NY, USA

^42^McLean Hospital, Belmont, MA, USA

^43^Division of Medical Sciences, Harvard Medical School, Boston, MA, USA

^44^Genomics Platform, Broad Institute of MIT and Harvard, Cambridge, MA, USA

^45^Department of Medicine, University of Mississippi Medical Center, Jackson, MI, USA

^46^Department of Epidemiology Colorado School of Public Health Aurora, CO, USA

^47^Institute for Molecular Medicine Finland, (FIMM) Helsinki, Finland

^48^Department of Medicine and Pharmacology, University of Illinois at Chicago, Chicago, IL, USA

^49^Vanderbilt University Medical Center, Nashville, TN, USA

^50^Department of Genetics, Texas Biomedical Research Institute, San Antonio, TX, USA

^51^Department of Biostatistics, Boston University School of Public Health, Boston, MA, USA

^52^National Heart Lung and Blood Institute’s Framingham Heart Study, Framingham, MA, USA

^53^Cardiac Arrhythmia Service and Cardiovascular Research Center, Massachusetts General Hospital, Boston, MA, USA

^54^Cardiovascular Epidemiology and Genetics Hospital del Mar Medical Research Institute, (IMIM) Barcelona Catalonia, Spain

^55^CIBER CV Barcelona, Catalonia, Spain

^56^Departament of Medicine, Medical School University of Vic-Central, University of Catalonia, Vic Catalonia, Spain

^57^Institute for Cardiogenetics, University of Lübeck, Lübeck, Germany

^58^German Research Centre for Cardiovascular Research, Hamburg/Lübeck/Kiel, Lübeck, Germany

^59^University Heart Center Lübeck, Lübeck, Germany

^60^Estonian Genome Center, Institute of Genomics University of Tartu, Tartu, Estonia

^61^Victor Chang Cardiac Research Institute, Darlinghurst, NSW, Australia

^62^Faculty of Medicine, UNSW Sydney, Kensington, NSW, Australia

^63^Cardiology Department, St Vincent’s Hospital, Darlinghurst, NSW, Australia

^64^Broad Genomics, Broad Institute of MIT and Harvard, Cambridge, MA, USA

^65^Diabetes Unit and Center for Genomic Medicine, Massachusetts General Hospital, Boston, MA, USA

^66^Programs in Metabolism and Medical & Population Genetics, Broad Institute of MIT and Harvard, Cambridge, MA, USA

^67^Institute of Clinical Molecular Biology, (IKMB) Christian-Albrechts-University of Kiel, Kiel, Germany

^68^Helsinki University and Helsinki University Hospital Clinic of Gastroenterology, Helsinki, Finland

^69^Bioinformatics Program MGH Cancer Center and Department of Pathology, Boston, MA, USA

^70^Cancer Genome Computational Analysis, Broad Institute of MIT and Harvard, Cambridge, MA, USA

^71^Department of Psychiatry and Behavioral Sciences, Boston Children’s Hospitaland Harvard Medical School, Boston, MA, USA

^72^Harvard Medical School Teaching Hospital, Boston, MA, USA

^73^Department of Endocrinology and Metabolism, Hadassah Medical Center and Faculty of Medicine, Hebrew University of Jerusalem, Israel

^74^Department of Psychiatry and Behavioral Sciences, SUNY Upstate Medical University, Syracuse, NY, USA

^75^Institute for Genomic Medicine, Columbia University Medical Center Hammer Health Sciences, New York, NY, USA

^76^Department of Genetics & Development Columbia University Medical Center, Hammer Health Sciences, New York, NY, USA

^77^Centro de Investigacion en Salud Poblacional, Instituto Nacional de Salud Publica, Mexico

^78^Lund University Sweden, Sweden

^79^Institute for Molecular Medicine Finland, (FIMM) HiLIFE University of Helsinki, Helsinki, Finland

^80^Division of Genetics and Genomics, Boston Children’s Hospital, Boston, MA, USA

^81^Lund University Diabetes Centre, Malmö, Skåne County, Sweden

^82^Washington School of Medicine, St Louis, MI, USA

^83^Human Genetics Center University of Texas Health Science Center at Houston, Houston, TX, USA

^84^Department of Neurology Columbia University, New York City, NY, USA

^85^Institute of Genomic Medicine, Columbia University, New York City, NY, USA

^86^Institute of Biomedicine, University of Eastern Finland, Kuopio, Finland

^87^Department of Psychiatry, Helsinki University Central Hospital Lapinlahdentie, Helsinki, Finland

^88^Department of Medical Epidemiology and Biostatistics, Karolinska Institutet, Stockholm, Sweden

^89^Icahn School of Medicine at Mount Sinai, New York, NY, USA

^90^Bonei Olam, Center for Rare Jewish Genetic Diseases, Brooklyn, NY, USA

^91^Department of Neurology, Helsinki University, Central Hospital, Helsinki, Finland

^92^Cardiovascular Disease Initiative and Program in Medical and Population Genetics, Broad Institute of MIT and Harvard, Cambridge, MA, USA

^93^Charles Bronfman Institute for Personalized Medicine, New York, NY, USA

^94^Division of Genome Science, Department of Precision Medicine, National Institute of Health, Republic of Korea

^95^MRC Centre for Neuropsychiatric Genetics & Genomics, Cardiff University School of Medicine, Cardiff, Wales

^96^Imperial College, Healthcare NHS Trust, London, UK

^97^National Heart and Lung Institute Cardiovascular Sciences, Hammersmith Campus, Imperial College London, London, UK

^98^Department of Health THL-National Institute for Health and Welfare, Helsinki, Finland

^99^Section of Cardiovascular Medicine, Department of Internal Medicine, Yale School of Medicine, Center for Outcomes Research and Evaluation Yale-New Haven Hospital, New Haven, CT, USA

^100^Division of Pediatric Gastroenterology, Emory University School of Medicine, Atlanta, GA, USA

^101^Department of Internal Medicine, Seoul National University Hospital, Seoul, Republic of Korea

^102^The University of Eastern Finland, Institute of Clinical Medicine, Kuopio, Finland

^103^Kuopio University Hospital, Kuopio, Finland

^104^Department of Genetics, Yale School of Medicine, New Haven, CT, USA

^105^Department of Clinical Chemistry Fimlab Laboratories and Finnish Cardiovascular Research Center-Tampere Faculty of Medicine and Health Technology, Tampere University, Finland

^106^The Mindich Child Health and Development, Institute Icahn School of Medicine at Mount Sinai, New York, NY, USA

^107^National Autonomous University of Mexico, Mexico City, Mexico

^108^Salvador Zubirán National Institute of Health Sciences and Nutrition, Mexico City, Mexico

^109^Li Ka Shing Institute of Health Sciences, The Chinese University of Hong Kong, Hong Kong, China

^110^Hong Kong Institute of Diabetes and Obesity, The Chinese University of Hong Kong, Hong Kong, China

^111^Centre for Population Genomics, Garvan Institute of Medical Research and UNSW Sydney, Sydney, Australia

^112^Centre for Population Genomics, Murdoch Children’s Research Institute, Melbourne, Australia

^113^University of California San Francisco Parnassus Campus, San Francisco, CA, USA

^114^Cardiovascular Research REGICOR Group, Hospital del Mar Medical Research Institute, (IMIM) Barcelona, Catalonia, Spain

^115^Department of Genetics, Harvard Medical School, Boston, MA, USA

^116^Oxford Centre for Diabetes, Endocrinology and Metabolism, University of Oxford, Churchill Hospital Old Road Headington, Oxford, OX, LJ, UK

^117^Welcome Centre for Human Genetics, University of Oxford, Oxford, OX, BN, UK

^118^Oxford NIHR Biomedical Research Centre, Oxford University Hospitals, NHS Foundation Trust, John Radcliffe Hospital, Oxford, OX, DU, UK

^119^John P. Hussman Institute for Human Genomics, Leonard M. Miller School of Medicine, University of Miami, Miami, FL, USA

^120^The Dr. John T. Macdonald Foundation Department of Human Genetics, Leonard M. Miller School of Medicine, University of Miami, Miami, FL, USA

^121^F. Widjaja Foundation Inflammatory Bowel and Immunobiology Research Institute Cedars-Sinai Medical Center, Los Angeles, CA, USA

^122^Atherogenomics Laboratory University of Ottawa, Heart Institute, Ottawa, Canada

^123^Division of General Internal Medicine, Massachusetts General Hospital, Boston, MA, USA

^124^Department of Clinical Sciences University, Hospital Malmo Clinical Research Center, Lund University, Malmö, Sweden

^125^Estonian Genome Center, Institute of Genomics, University of Tartu, Tartu, Estonia

^126^University of Arizona Health Science, Tuscon, AZ, USA

^127^Department of Medicine, University of Maryland School of Medicine, Baltimore, MD, USA

^128^Howard Hughes Medical Institute and Department of Molecular Biology, Massachusetts General Hospital, Boston, MA, USA

^129^International Centre for Diarrhoeal Disease Research, Bangladesh

^130^Perelman School of Medicine, University of Pennsylvania, Philadelphia, PA, USA

^131^Johns Hopkins Bloomberg School of Public Health, Baltimore, MD, USA

^132^Lund University, Dept. Clinical Sciences, Skåne University Hospital, Malmö, Sweden

^133^Department of Statistical Genetics, Osaka University Graduate School of Medicine, Suita, Japan

^134^Laboratory of Statistical Immunology, Immunology Frontier Research Center (WPI-IFReC), Osaka University, Suita, Japan

^135^Integrated Frontier Research for Medical Science Division, Institute for Open and Transdisciplinary Research Initiatives, Osaka University, Suita, Japan

^136^Instituto Nacional de Medicina Genómica, (INMEGEN) Mexico City, Mexico

^137^Medical Research Institute, Ninewells Hospital and Medical School University of Dundee, Dundee, UK

^138^Wake Forest School of Medicine, Winston-Salem, NC, USA

^139^Department of Molecular Medicine and Biopharmaceutical Sciences, Graduate School of Convergence Science and Technology, Seoul National University, Seoul, Republic of Korea

^140^Department of Psychiatry Keck School of Medicine at the University of Southern California, Los Angeles, CA, USA

^141^Department of Psychiatry and Behavioral Sciences, Johns Hopkins University School of Medicine, Baltimore, MD, USA

^142^Children’s Hospital of Philadelphia, Philadelphia, PA, USA

^143^Division of Genetics and Epidemiology, Institute of Cancer Research, London, SM, NG

^144^University of Washington, Seattle, WA, USA

^145^Fred Hutchinson Cancer Research Center, Seattle, WA, USA

^146^Medical Research Center, Oulu University Hospital, Oulu Finland and Research Unit of Clinical Neuroscience Neurology University of Oulu, Oulu, Finland

^147^Center for Public Health Genomics, University of Virginia, Charlottesville, VA, USA

^148^Department of Public Health Sciences, University of Virginia, Charlottesville, VA, USA

^149^Research Center Montreal Heart Institute, Montreal, Quebec, Canada

^150^Department of Medicine, Faculty of Medicine Université de Montréal, Québec, Canada

^151^Department of Public Health Faculty of Medicine, University of Helsinki, Helsinki, Finland

^152^Broad Institute of MIT and Harvard, Cambridge, MA, USA

^153^Department of Biomedical Informatics Vanderbilt, University Medical Center, Nashville, TN, USA

^154^Department of Medicine, Vanderbilt University Medical Center, Nashville, TN, USA

^155^The Institute for Translational Genomics and Population Sciences, Department of Pediatrics, The Lundquist Institute for Biomedical Innovation at Harbor-UCLA Medical Center, Torrance, CA, USA

^156^Department of Biostatistics and Epidemiology, Perelman School of Medicine, University of Pennsylvania, Philadelphia, PA, USA

^157^Department of Medicine, Perelman School of Medicine at the University of Pennsylvania, Philadelphia, PA, USA

^158^Center for Non-Communicable Diseases, Karachi, Pakistan

^159^National Institute for Health and Welfare, Helsinki, Finland

^160^Deutsches Herzzentrum, München, Germany

^161^Technische Universität München, Germany

^162^Institute of Genetic Epidemiology, Department of Genetics and Pharmacology, Medical University of Innsbruck, 6020 Innsbruck, Austria

^163^Duke Molecular Physiology Institute, Durham, NC

^164^Division of Cardiovascular Medicine, Nashville VA Medical Center, Vanderbilt University School of Medicine, Nashville, TN, USA

^165^Division of Endocrinology, National University Hospital, Singapore

^166^NUS Saw Swee Hock School of Public Health, Singapore

^167^Channing Division of Network Medicine, Brigham and Women’s Hospital, Boston, MA, USA

^168^Harvard Medical School, Boston, MA, USA

^169^Department of Psychiatry, Icahn School of Medicine at Mount Sinai, New York, NY, USA

^170^Department of Genetics and Genomic Sciences, Icahn School of Medicine at Mount Sinai, New York, NY, USA

^171^Institute for Genomics and Multiscale Biology, Icahn School of Medicine at Mount Sinai, New York, NY, USA

^172^The Wallenberg Laboratory/Department of Molecular and Clinical Medicine, Institute of Medicine, Gothenburg University and the Department of Cardiology, Sahlgrenska University Hospital, Gothenburg, Sweden

^173^Department of Cardiology, Wallenberg Center for Molecular Medicine and Lund University Diabetes Center, Clinical Sciences, Lund University and Skåne University Hospital, Lund, Sweden

^174^Institute of Clinical Medicine Neurology, University of Eastern Finad, Kuopio, Finland

^175^Sorbonne Université, INSERM, Centre de Recherche Saint-Antoine, CRSA, AP-HP, Saint Antoine Hospital, Gastroenterology department, F-75012 Paris, France

^176^INRA, UMR1319 Micalis & AgroParisTech, Jouy en Josas, France

^177^Paris Center for Microbiome Medicine, (PaCeMM) FHU, Paris, France

^178^Department of Twin Research and Genetic Epidemiology King’s College London, London, UK

^179^The McDonnell Genome Institute at Washington University, Seattle, WA, USA

^180^Departments of Genetics and Psychiatry, University of North Carolina, Chapel Hill, NC, USA

^181^Saw Swee Hock School of Public Health National University of Singapore, National University Health System, Singapore

^182^Department of Medicine, Yong Loo Lin School of Medicine National University of Singapore, Singapore

^183^Duke-NUS Graduate Medical School, Singapore

^184^Life Sciences Institute, National University of Singapore, Singapore

^185^Department of Statistics and Applied Probability, National University of Singapore, Singapore

^186^Center for Behavioral Genomics, Department of Psychiatry, University of California, San Diego, CA, USA

^187^Institute of Genomic Medicine, University of California San Diego, San Diego, CA, USA

^188^Endocrinology, Abdominal Center, Helsinki University Hospital, Helsinki, Finland

^189^Institute of Genetics, Folkhalsan Research Center, Helsinki, Finland

^190^Juliet Keidan Institute of Pediatric Gastroenterology Shaare Zedek Medical Center, The Hebrew University of Jerusalem, Jerusalem, Israel

^191^Instituto de Investigaciones Biomédicas, UNAM, Mexico City, Mexico

^192^Instituto Nacional de Ciencias Médicas y Nutrición Salvador Zubirán, Mexico City, Mexico

^193^Department of Public Health Faculty of Medicine University of Helsinki, Helsinki, Finland

^194^Department of Psychiatry and Human Behavior, University of California Irvine, Irvine, CA, USA

^195^National Heart & Lung Institute & MRC London Institute of Medical Sciences, Imperial College, London, UK

^196^Royal Brompton & Harefield Hospitals, Guy’s and St. Thomas’ NHS Foundation Trust, London, UK

^197^Radcliffe Department of Medicine, University of Oxford, Oxford, UK

^198^Department of Gastroenterology and Hepatology, University of Groningen and University Medical Center Groningen, Groningen, Netherlands

^199^Folkhälsan Institute of Genetics, Folkhälsan Research Center, Helsinki, Finland

^200^National Heart & Lung Institute and MRC London Institute of Medical Sciences, Imperial College London, London, UK

^201^Cardiovascular Research Centre, Royal Brompton & Harefield Hospitals NHS Trust, London, UK

^202^Department of Physiology and Biophysics, University of Mississippi Medical Center, Jackson, MS, USA

^203^Program in Infectious Disease and Microbiome, Broad Institute of MIT and Harvard, Cambridge, MA, USA

^204^Center for Computational and Integrative Biology, Massachusetts General Hospital, Boston, MA, USA

Authors received funding as follows:

Emelia J Benjamin: Framingham Heart Study 75N92019D00031; and R01HL092577

Matthew J. Bown: British Heart Foundation awards CS/14/2/30841 and RG/18/10/33842

Josée Dupuis: National Heart Lung and Blood Institute’s Framingham Heart Study Contract (HHSNI); National Institute for Diabetes and Digestive and Kidney Diseases (NIDDK) R DK

Martti Färkkilä: State funding for university level health research Laura D. Gauthier: Intel, Illumina

Stephen J. Glatt: U.S. NIMH Grant R MH

Leif Groop: The Academy of Finland and University of Helsinki: Center of Excellence for Complex Disease Genetics (grant number 312063 and 336822), Sigrid Jusélius Foundation; IMI 2 (grant No 115974 and 15881)

Mikko Hiltunen: Academy of Finland (grant 338182) Sigrid Jusélius Foundation the Strategic Neuroscience Funding of the University of Eastern Finland

Chaim Jalas: Bonei Olam

Jaakko Kaprio: Academy of Finland (grants 312073 and 336823)

Jacob McCauley: National Institute of Diabetes and Digestive and Kidney Disease Grant R01DK104844

Yukinori Okada: JSPS KAKENHI (19H01021, 20K21834), AMED (JP21km0405211, JP21ek0109413, JP21gm4010006, JP21km0405217, JP21ek0410075), JST Moonshot R&D (JPMJMS2021)

Michael J. Owen: Medical Research Council UK: Centre Grant No. MR/L010305/1, Program Grant No. G0800509

Aarno Palotie: the Academy of Finland Center of Excellence for Complex Disease Genetics (grant numbers 312074 and 336824) and Sigrid Jusélius Foundation

John D. Rioux: National Institute of Diabetes and Digestive and Kidney Diseases (NIDDK; DK062432), from the Canadian Institutes of Health (CIHR GPG 102170), from Genome Canada/Génome Québec (GPH-129341), and a Canada Research Chair (#230625)

Samuli Ripatti: the Academy of Finland Center of Excellence for Complex Disease Genetics (grant number) Sigrid Jusélius Foundation

Jerome I. Rotter: Trans-Omics in Precision Medicine (TOPMed) program was supported by the National Heart, Lung and Blood Institute (NHLBI). WGS for “NHLBI TOPMed: Multi-Ethnic Study of Atherosclerosis (MESA)” (phs001416.v1.p1) was performed at the Broad Institute of MIT and Harvard (3U54HG003067-13S1). Core support including centralized genomic read mapping and genotype calling, along with variant quality metrics and filtering were provided by the TOPMed Informatics Research Center (3R01HL-117626-02S1; contract HHSN268201800002I). Core support including phenotype harmonization, data management, sample-identity QC, and general program coordination were provided by the TOPMed Data Coordinating Center (R01HL-120393; U01HL-120393; contract HHSN268201800001I). We gratefully acknowledge the studies and participants who provided biological samples and data for MESA and TOPMed. JSK was supported by the Pulmonary Fibrosis Foundation Scholars Award and grant K23-HL-150301 from the NHLBI. MRA was supported by grant K23-HL-150280, AJP was supported by grant K23-HL-140199, and AM was supported by R01-HL131565 from the NHLBI. EJB was supported by grant K23-AR-075112 from the National Institute of Arthritis and Musculoskeletal and Skin Diseases.The MESA project is conducted and supported by the National Heart, Lung, and Blood Institute (NHLBI) in collaboration with MESA investigators. Support for MESA is provided by contracts 75N92020D00001, HHSN268201500003I, N01-HC-95159, 75N92020D00005, N01-HC-95160, 75N92020D00002, N01-HC-95161, 75N92020D00003, N01-HC-95162, 75N92020D00006, N01-HC-95163, 75N92020D00004, N01-HC-95164, 75N92020D00007, N01-HC-95165, N01-HC-95166, N01-HC-95167, N01-HC-95168, N01-HC-95169, UL1-TR-000040, UL1-TR-001079, and UL1-TR-001420. Also supported in part by the National Center for Advancing Translational Sciences, CTSI grant UL1TR001881, and the National Institute of Diabetes and Digestive and Kidney Disease Diabetes Research Center (DRC) grant DK063491 to the Southern California Diabetes Endocrinology Research Center

Edwin K. Silverman: NIH Grants U01 HL089856 and U01 HL089897

J. Gustav Smith: The Swedish Heart-Lung Foundation (2019-0526), the Swedish Research Council (2017-02554), the European Research Council (ERC-STG-2015-679242), Skåne University Hospital, governmental funding of clinical research within the Swedish National Health Service, a generous donation from the Knut and Alice Wallenberg foundation to the Wallenberg Center for Molecular Medicine in Lund, and funding from the Swedish Research Council (Linnaeus grant Dnr 349-2006-237, Strategic Research Area Exodiab Dnr 2009-1039) and Swedish Foundation for Strategic Research (Dnr IRC15-0067) to the Lund University Diabetes Center

Kent D. Taylor: Trans-Omics in Precision Medicine (TOPMed) program was supported by the National Heart, Lung and Blood Institute (NHLBI). WGS for “NHLBI TOPMed: Multi-Ethnic Study of Atherosclerosis (MESA)” (phs001416.v1.p1) was performed at the Broad Institute of MIT and Harvard (3U54HG003067-13S1). Core support including centralized genomic read mapping and genotype calling, along with variant quality metrics and filtering were provided by the TOPMed Informatics Research Center (3R01HL-117626-02S1; contract HHSN268201800002I). Core support including phenotype harmonization, data management, sample-identity QC, and general program coordination were provided by the TOPMed Data Coordinating Center (R01HL-120393; U01HL-120393; contract HHSN268201800001I). We gratefully acknowledge the studies and participants who provided biological samples and data for MESA and TOPMed. JSK was supported by the Pulmonary Fibrosis Foundation Scholars Award and grant K23-HL-150301 from the NHLBI. MRA was supported by grant K23-HL-150280, AJP was supported by grant K23-HL-140199, and AM was supported by R01-HL131565 from the NHLBI. EJB was supported by grant K23-AR-075112 from the National Institute of Arthritis and Musculoskeletal and Skin Diseases.The MESA project is conducted and supported by the National Heart, Lung, and Blood Institute (NHLBI) in collaboration with MESA investigators. Support for MESA is provided by contracts 75N92020D00001, HHSN268201500003I, N01-HC-95159, 75N92020D00005, N01-HC-95160, 75N92020D00002, N01-HC-95161, 75N92020D00003, N01-HC-95162, 75N92020D00006, N01-HC-95163, 75N92020D00004, N01-HC-95164, 75N92020D00007, N01-HC-95165, N01-HC-95166, N01-HC-95167, N01-HC-95168, N01-HC-95169, UL1-TR-000040, UL1-TR-001079, and UL1-TR-001420. Also supported in part by the National Center for Advancing Translational Sciences, CTSI grant UL1TR001881, and the National Institute of Diabetes and Digestive and Kidney Disease Diabetes Research Center (DRC) grant DK063491 to the Southern California Diabetes Endocrinology Research Center

Tiinamaija Tuomi: The Academy of Finland and University of Helsinki: Center of Excellence for Complex Disease Genetics (grant number 312072 and 336826), Folkhalsan Research Foundation, Helsinki University Hospital, Ollqvist Foundation, Liv och Halsa foundation; NovoNordisk Foundation

Teresa Tusie-Luna: CONACyT Project 312688

James S. Ware: Sir Jules Thorn Charitable Trust [21JTA], Wellcome Trust [107469/Z/15/Z], Medical Research Council (UK), NIHR Imperial College Biomedical Research Centre

Rinse K. Weersma: The Lifelines Biobank initiative has been made possible by subsidy from the Dutch Ministry of Health Welfare and Sport the Dutch Ministry of Economic Affairs the University Medical Centre Groningen (UMCG the Netherlands) the University of Groningen and the Northern Provinces of the Netherlands

No conflicts of interest to declare.

